# Plasma membrane-associated graphene oxide as a platform for modulating signalling through cell-surface receptors: an integrin-focused proof-of-concept study

**DOI:** 10.64898/2026.08.25.747094

**Authors:** Angeliki Karakasidi, Neus Lozano, Kostas Kostarelos, Sandra Vranic

## Abstract

Graphene oxide (GO) has primarily been investigated as a carrier for intracellular delivery of therapeutic molecules. In previous work, we identified a cell type-dependent interaction pattern in which GO remained predominantly associated with the plasma membrane of cancer cells but was internalised by non-cancerous epithelial cells. Here, we explored whether plasma membrane-associated GO can be used as a platform to present bioactive ligands and influence cell-surface receptor signalling in cancer cells. To test this hypothesis, we targeted integrin receptors at the plasma membrane in glioblastoma cell models using an RGD-containing peptide non-covalently complexed with GO. We assessed GO-peptide interactions, cellular interactions/uptake, motility, and focal adhesion signalling readouts.

Peptide association was quantified using a 2,4,6-trinitrobenzene sulfonic acid (TNBSA) assay, and GO was characterised by atomic force microscopy, X-ray photoelectron spectroscopy, X-ray diffraction, and colloidal measurements. Immediately after complexation, ∼70% of RGD was associated with GO. Peptide association increased the nitrogen signal and shifted the principal GO XRD peak while retaining nanosheet morphology. Biological responses were examined in U87 and U251 glioblastoma cells with different integrin-positive fractions, and in non-cancerous BEAS-2B bronchial epithelial cells. Confocal microscopy showed that GO and GO:RGD remained predominantly localised on the plasma membrane in U87 and U251 cells, whereas greater intracellular localisation was observed in BEAS-2B cells.

Importantly, GO:RGD significantly reduced key indicators of cell motility: cell velocity in U87 and U251 cells, with trajectory and mean-square-displacement analyses supporting restricted cellular movement. Free RGD had no significant effect, while GO alone produced a smaller reduction in motility only in U251 cells. No treatment significantly altered BEAS-2B motility. Flow cytometry also showed a reduced pFAK-associated signal in GO:RGD-treated U87 cells.

These findings establish a proof of concept that the cell-line-dependent plasma membrane localisation of GO can be exploited as a membrane-associated nano-bio interface for cell-surface-active ligands, opening the way for the development of GO-based platforms that modulate receptor-mediated signalling and cell behaviour.

## Introduction

Graphene oxide (GO), the oxidised derivative of graphene, has attracted considerable interest in nanomedicine [1,2]. Its high surface area and oxygen-containing groups, including carboxyl, hydroxyl and epoxy groups, enable covalent functionalisation as well as non-covalent interactions through electrostatic forces, hydrogen bonding, van der Waals forces and π–π stacking [3,4]. The coexistence of carbon domains and oxygen-rich regions also gives GO an amphiphilic character that supports its dispersion in aqueous environments. Together, these properties have established GO as a versatile nanocarrier, particularly for drug and biomolecule delivery [5,6].

The development of GO-based biomedical applications requires a detailed understanding of how GO interacts with cells [7]. In our earlier work, Chen et al. showed that thin GO nanosheets entered non-phagocytic BEAS-2B cells predominantly through macropinocytosis [8]. Our subsequent comparative analysis revealed a distinctive interaction pattern between GO and cancer and non-cancer cells [9]. Specifically, GO was internalised by non-cancerous epithelial cells, whereas it remained predominantly associated with the plasma membrane of the cancer cell lines examined. This distinctive membrane association provides a strong rationale for exploiting GO as a platform to deliver plasma-membrane-targeting therapeutic molecules to the cancer-cell surface.

Cell-surface receptors regulate processes central to cancer progression. Among these, integrins are transmembrane receptors that connect the extracellular matrix to the cytoskeleton and transmit bidirectional signals across the plasma membrane [10,11]. Dysregulated integrin expression and signalling promote cancer-cell adhesion, migration, invasion and resistance to therapy. In glioblastoma, an aggressive and highly invasive primary brain tumour, the RGD-binding integrins αvβ3 and αvβ5 contribute to tumour invasion and angiogenesis, whereas α5β1 is implicated in tumour progression and resistance to therapy [12–15]. Functional inhibition of these receptors can therefore disrupt tumour-cell adhesion and motility and interfere with pro-survival signalling pathways, making integrins promising targets for therapeutic intervention.

On this basis, we hypothesised that the preferential association of GO with the plasma membrane of cancer cells could be harnessed to modulate cell-surface receptor function. To examine this hypothesis, we selected integrins as a model receptor system and employed a ligand containing the Arg-Gly-Asp (RGD) sequence, which is recognised by multiple integrin subtypes and can inhibit integrin-dependent cellular functions [16,17]. While nanomaterials have frequently been used to present RGD ligands to enhance receptor-mediated uptake, such strategies prioritise internalisation and intracellular delivery [18–20]. In contrast, maintaining ligand availability at the plasma membrane may be more advantageous when targeting cell-surface receptors [21]. Plasma-membrane-directed nanodelivery has also been explored for therapeutic agents whose activity depends on extracellular receptor engagement, including the presentation or release of TRAIL at cancer-cell membranes using single-walled carbon nanotubes, graphene-based sequential-delivery systems and transformable DNA nanocarriers [22–24]. However, these systems rely on engineered ligand presentation or stimulus-responsive release, whereas exploiting the intrinsic cell-dependent membrane association of a nanomaterial could provide a distinct approach for sustaining ligand-receptor interactions at the cell surface.

As a proof of concept, GO was non-covalently complexed with the linear RGD-containing peptide GRGDS, and peptide association and the physicochemical properties of the resulting GO:RGD formulation were fully characterised. Biological responses to GO:RGD were examined in U87 and U251 glioblastoma cells, which display different integrin profiles, and in BEAS-2B bronchial epithelial cells as a non-cancerous comparator. We evaluated GO:RGD localisation relative to the plasma membrane and αvβ3-labelled membrane regions, together with its effects on single-cell motility and FAK Tyr397 phosphorylation as a downstream readout of integrin-linked signalling [25]. Collectively, these experiments tested whether the GO:RGD formulation retained the membrane-associated behaviour of GO alone and modulated integrin-linked motility and signalling in relation to cell-specific integrin profiles. By examining GO as a membrane-associated platform, this study extends its conventional use and explores its potential as a nano-bio interface for delivering receptor-targeting ligands and modulating signalling at the cell surface.

## Results

### Preparation and physicochemical characterisation of the GO:RGD formulation

Detailed physicochemical characterisation of the starting GO material is reported in Table S1; briefly, the majority of nanosheets (95%) had lateral dimension below 850 nm and 1-2 nm thickness. The RGD peptide was non-covalently complexed with GO as shown in Figure 1A. GO and RGD were used at concentrations of 25 µg/mL and 5 µg/mL, respectively, corresponding to a 10:2 GO:RGD mass ratio, and mixed under aqueous conditions for 30 min at room temperature. Following complexation, the GO:RGD preparation was subjected to four sequential centrifugal-filtration washes (2,000 × g for 5 min; Amicon Ultra-4 Centrifugal Filter Unit, 100 kDa, 4 mL; EMD Millipore, UK) to remove unbound RGD. Dispersion behaviour was evaluated by visual observation (Figure 1B). Representative images of GO, purified GO:RGD and RGD were acquired at 0, 4 and 24 h (Figure 1B). No visible sedimentation was observed for GO or GO:RGD during this period. Following removal of unbound peptide, the pH and colloidal stability parameters of GO and GO:RGD were monitored at the same time points (Figure 1C). The pH of GO and GO:RGD remained approximately 3.7, whereas that of the RGD control remained approximately 4.5. DLS size, PDI and zeta potential showed little variation over 24 h. At 24 h, both GO and GO:RGD had DLS size of ∼200 nm. The corresponding zeta potential and PDI values were −45 mV and 0.28 for GO:RGD and −52 mV and 0.21 for GO, respectively. The intensity-weighted size distributions are shown in Figure S1 confirming the findings.

**Figure 1.**
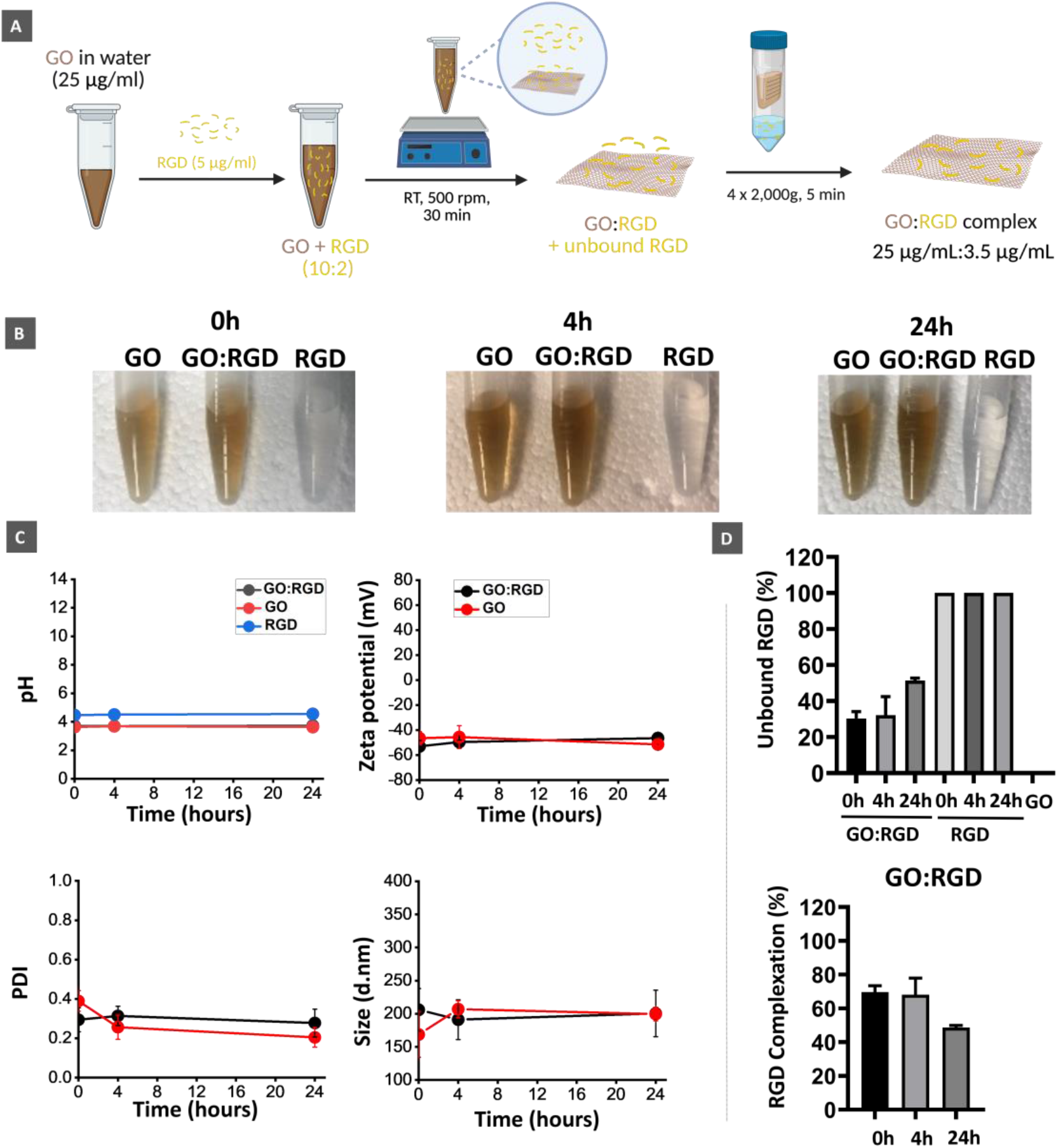
Complexation and physicochemical characterisation of GO:RGD. (A) Schematic of the non-covalent complexation of GO with RGD at a 10:2 GO:RGD mass ratio and the analytical purification used to remove the unbound peptide. At an initial concentration of 25 µg/mL GO and 5 µg/mL RGD, the 69.7% associated fraction corresponded to approximately 3.5 µg/mL RGD after purification. (B) Representative images of GO, purified GO:RGD and RGD preparations at 0, 4 and 24 h. (C) pH, DLS size, PDI and zeta potential of the indicated complexations. Data are presented as mean ± SD from three independently prepared samples, each measured in technical triplicate. (D) TNBSA-derived percentages of unbound and calculated GO-associated RGD at 0, 4 and 24 h. Data are presented as mean ± SD from three independent experiments.

Peptide association with GO was quantified using the TNBSA assay (Figure 1D). TNBSA analysis was used to quantify unbound peptide and calculate the fraction associated with GO at 0, 4 and 24 h (Figure 1D). Immediately after complexation, 30.3% of the total RGD was detected as unbound, corresponding to a calculated GO-associated fraction of 69.7%. At 4 h, the unbound and associated fractions were 32.5% and 67.5%, respectively. At 24 h, 51.2% of the peptide was detected as unbound and 48.8% remained associated with GO. At an initial RGD concentration of 5 µg/mL, the 69.7% associated fraction corresponded to approximately 3.5 µg/mL RGD following purification. The GO-only control produced only background-level signal. Absorbance values were measured in triplicate, averaged and converted into peptide concentrations using the RGD calibration curve shown in Figure S2. The corresponding values are reported in Tables S2 and S3.

### Structural and surface characterisation of the GO:RGD

The morphology of purified GO:RGD was compared with that of GO using AFM (Figure 2A). The AFM height images (Figure 2A, top) showed sheet-like structures for both preparations, with representative thicknesses of ∼1.5 nm, consistent with predominantly single-layer GO. The corresponding cross-sectional profiles (Figure 2A, bottom) showed minimal differences between GO and GO:RGD, indicating that the overall nanosheet morphology was retained following peptide association.

**Figure 2.**
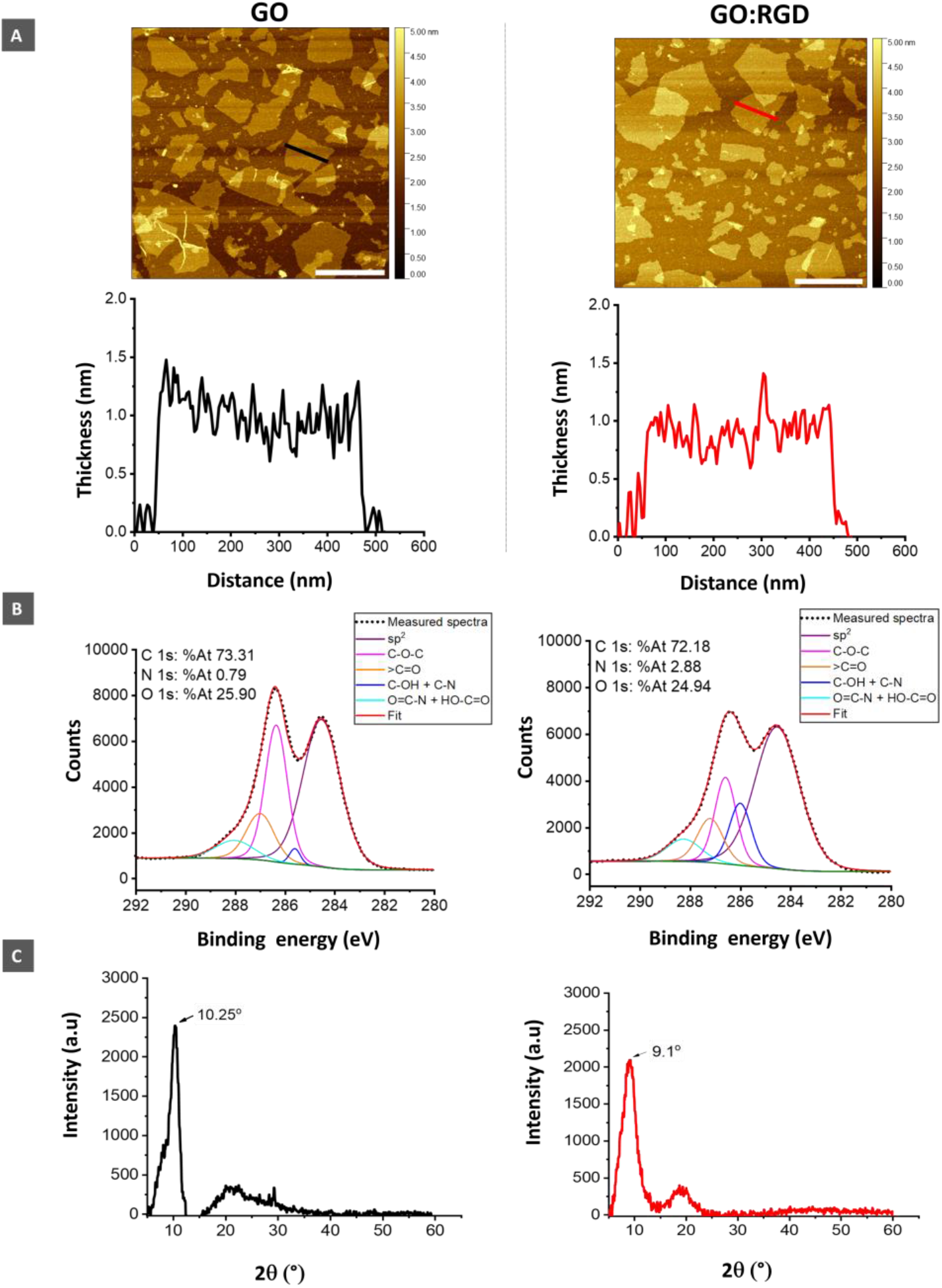
Structural and surface characterisation of GO and the purified GO:RGD complex. (A) Representative AFM height images and corresponding line profiles showing the sheet-like morphology and thickness of GO and GO:RGD. Scale bar = 500 nm. Distance is the length along the selected line. (B) High-resolution C 1s XPS spectra with fitted components and elemental atomic percentages, showing an increase in nitrogen from 0.79 % in GO to 2.88 % in GO:RGD, consistent with peptide association. (C) XRD patterns of dried GO and GO:RGD samples, showing a shift in the principal GO peak from 10.25° to 9.1° following peptide association.

Peptide association was further examined using XPS and XRD. XPS detected an increase in the nitrogen atomic fraction from approximately 0.8 % in GO to 2.88 % in GO:RGD (Figure 2B), consistent with the presence of the RGD-containing peptide in the purified complex. XRD analysis showed that the principal GO peak shifted to a lower 2θ value, from 10.25° for GO to 9.1° for GO:RGD (Figure 2C). This shift is consistent with increased interlayer spacing in the dried GO:RGD sample. Collectively, the AFM, XPS and XRD measurements support the association of RGD with GO while showing that the overall nanosheet morphology was retained.

### Biological effects of GO:RGD

#### Biocompatibility and interactions with the plasma membrane

Free RGD was evaluated over a concentration range of 0–100 µg/mL for 24 h in U87, U251 and BEAS-2B cells as an exploratory dose-selection experiment (Figure S3). At 100 µg/mL RGD, viability remained above 90% in U87 cells and above 80% in U251 and BEAS-2B cells (Figure S3B). Despite the preservation of viability, optical microscopy revealed concentration-dependent clustering of U87 cells from 25 µg/mL, which became more pronounced at higher concentrations. This morphology was not observed in U251 or BEAS-2B cells across the tested concentration range (Figure S3A). The U87 clustering was therefore considered a dose-limiting morphological response. On this basis, RGD concentration of 5 µg/mL was selected for confocal imaging and motility experiments, while 10 µg/mL was used for the pFAK experiment. The GO concentrations used here were selected based on previous studies showing their lack of cytotoxicity under comparable exposure conditions [9]. Consistent with this, GO:RGD-treated cells retained normal morphology during confocal and time-lapse imaging, with no overt evidence of treatment-related cytotoxicity.

The localisation of GO and GO:RGD relative to the plasma membrane was examined after 24 h using the intrinsic red fluorescence of GO and CellMask staining of the plasma membrane [8, 26]. Cells were untreated, RGD (5 µg/mL), GO (25 µg/mL) or GO:RGD (25:5 µg/mL) treated, and representative confocal images were examined using orthogonal projections (Figure 3). In U87 and U251 cells, the red signal from both GO and GO:RGD was predominantly associated with the cell periphery and occupied the same peripheral regions as the CellMask-labelled plasma membrane. In BEAS-2B cells, a greater proportion of the red signal was observed within the cells, consistent with greater intracellular localisation. These results confirm the cell-line-dependent localisation and indicate that association with RGD did not produce an obvious redistribution of GO from the cell periphery to the cell interior in either glioblastoma cell line.

**Figure 3.**
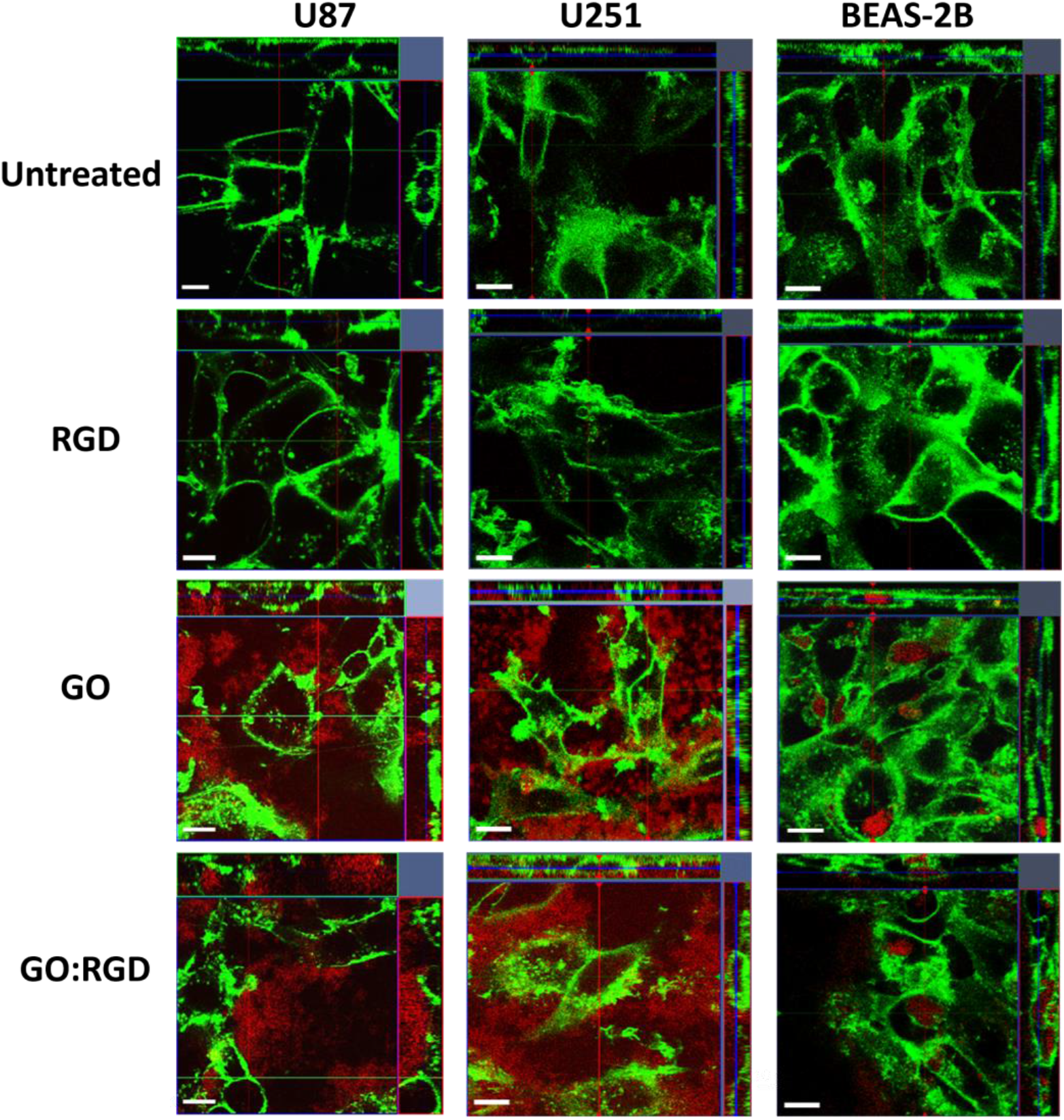
Cell-line-dependent localisation of GO and GO:RGD relative to the plasma membrane. U87, U251 and BEAS-2B cells were untreated, RGD (5 µg/mL), GO (25 µg/mL) or GO:RGD (25:5 µg/mL) treated for 24 h. The plasma membrane was labelled with CellMask Green (green), while GO and GO:RGD were visualised through the intrinsic fluorescence of GO (red). Representative central optical sections are shown together with the corresponding orthogonal projections. Scale bars, 10 µm.

#### Spatial association of GO:RGD with αvβ3-labelled membrane regions

Baseline integrin profiles were subsequently determined by flow cytometry (Figure S4). The αvβ3-, αvβ5- and α5β1-positive populations were approximately 90%, 65% and 95%, respectively, in U87 cells; 23%, 37% and 11% in U251 cells; and 15%, 23% and 21% in BEAS-2B cells. U87 therefore displayed the highest fractions of integrin-positive cells for all three integrins, whereas U251 and the non-cancerous bronchial epithelial comparator BEAS-2B displayed lower proportion of integrin-positive cells. These differences provided a basis for examining the localisation of GO relative to αvβ3-labelled membrane regions across the three cell lines.

To examine this spatial relationship, cells were untreated, RGD (5 µg/mL), GO (25 µg/mL) or GO:RGD (25:5 µg/mL) treated for 24 h, followed by αvβ3 immunofluorescence staining and confocal microscopy (Figure 4). In U87 and U251 cells, the red signal from both GO and GO:RGD was observed predominantly at the cell periphery and in the same membrane regions as αvβ3 labelling. In BEAS-2B cells, a greater proportion of the GO-associated red signal appeared within the cell interior, consistent with the cell-line-dependent localisation observed using plasma-membrane staining. Additional single-channel and merged confocal images for U87, U251 and BEAS-2B cells are provided in Figures S5–S7. Collectively, these images support spatial proximity between membrane-associated GO:RGD and αvβ3-labelled regions in the glioblastoma cells.

**Figure 4.**
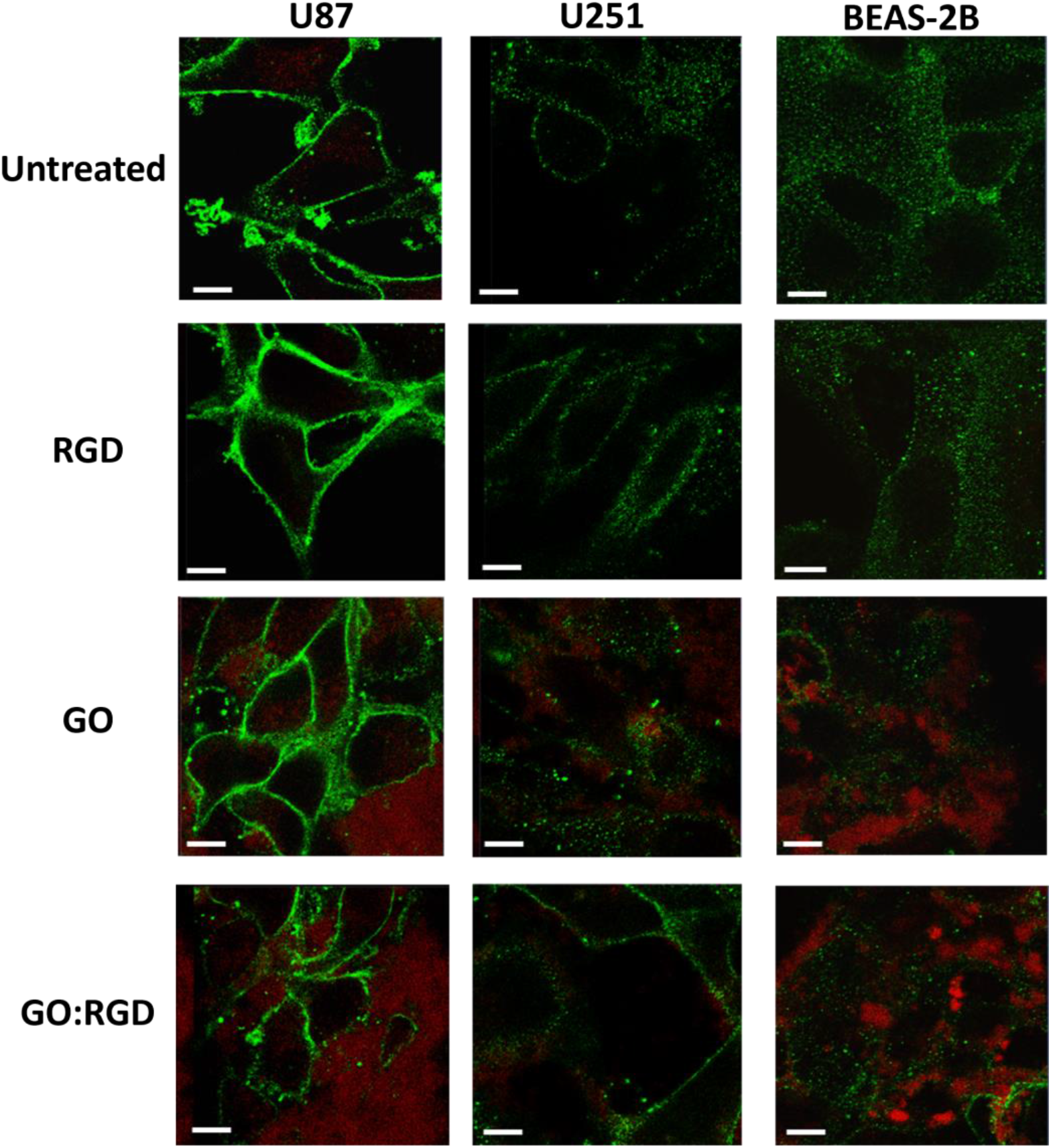
Spatial association of GO:RGD with αvβ3-positive membrane regions. U87, U251 and BEAS-2B cells were untreated, RGD (5 µg/mL), GO (25 µg/mL) or GO:RGD (25:5 µg/mL) treated for 24 h. αvβ3 was visualised by immunofluorescence staining (green), while GO and GO:RGD were visualised through the intrinsic fluorescence of GO (red). Representative merged confocal images are shown. Additional single-channel and merged images are provided in Figures S5–S7. Scale bars, 10 µm.

#### GO:RGD reduces mean cell velocity in glioblastoma cells

Single-cell motility was assessed by 24 h live-cell time-lapse microscopy in U87, U251 and BEAS-2B cells exposed to cell culture medium alone (untreated), RGD (5 µg/mL), GO (25 µg/mL) or GO:RGD (25:5 µg/mL). Mean cell velocity was used as the quantitative endpoint, while origin-centred trajectories and mean-square-displacement (MSD) profiles were used as descriptive representations of cell movement (Figures 5 and 6).

**Figure 5.**
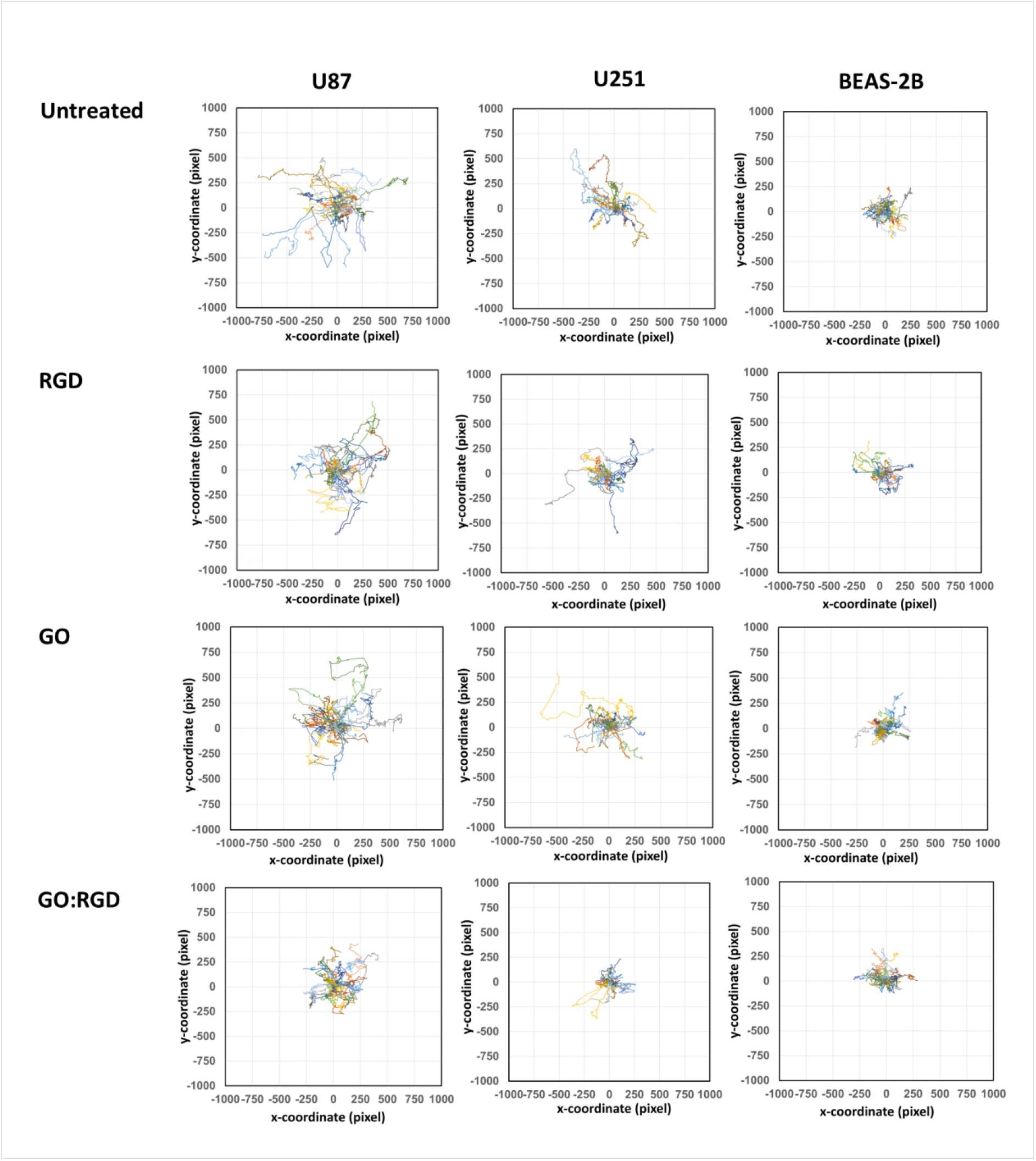
Origin-centred cell trajectories following GO:RGD exposure. Origin-centred trajectories of U87, U251 and BEAS-2B cells exposed to medium alone, RGD (5 µg/mL), GO (25 µg/mL) or GO:RGD (25:5 µg/mL) for 24 h. Individual trajectories were translated to a common origin to visualise cell-displacement patterns. Representative trajectories from 25 tracked cells per condition are shown. Twenty-five cells per condition were tracked in each of three independent experiments using CellTracker, and trajectories were plotted using DiPer.

**Figure 6.**
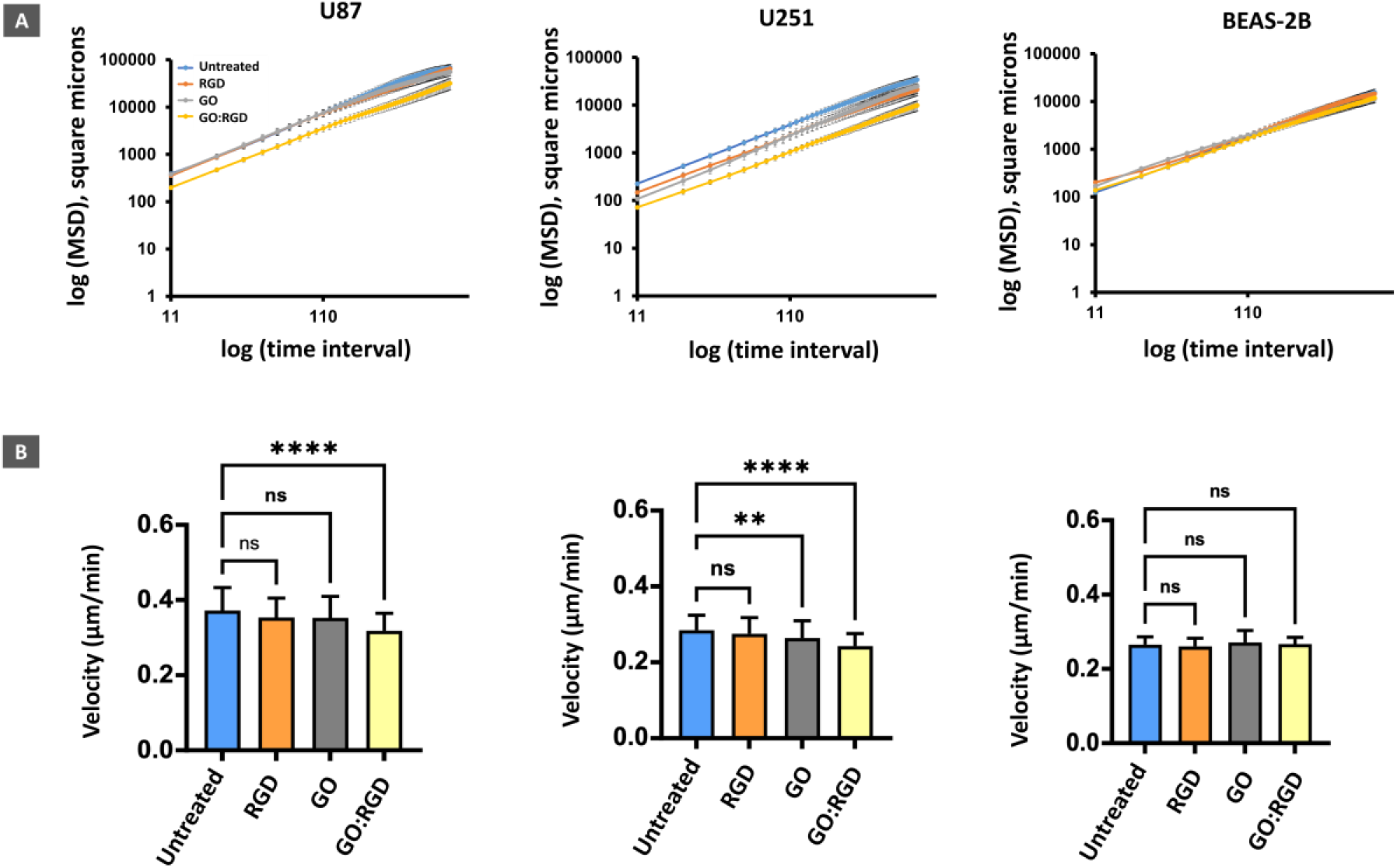
Mean-square-displacement and velocity profiles following treatments. (A) Mean-square displacement over time for U87, U251 and BEAS-2B cells exposed to medium alone, RGD (5 µg/mL), GO (25 µg/mL) or GO:RGD (25:5 µg/mL) for 24 h. MSD profiles were calculated from trajectories obtained using CellTracker and DiPer. (B) Mean cell velocity in U87, U251 and BEAS-2B cell lines following the indicated treatments. Statistical analysis was performed using one-way ANOVA, followed by Dunnett’s multiple-comparisons test against the untreated control, where *, **, ***, and **** indicate P<0.05, P<0.01, P<0.001, and P<0.0001, respectively. Data are presented as mean ± SEM of the three independent experiments (n = 3), with each experimental value representing the mean velocity of 25 cells.

Relative to untreated cells, GO:RGD significantly reduced mean velocity in both U87 and U251 cells (*p* < 0.0001; Figure 6B). Free RGD did not significantly alter mean velocity in either glioblastoma cell line. GO alone did not significantly affect U87 velocity but produced a smaller, statistically significant reduction in U251 cells (*p* < 0.01). No treatment significantly altered mean velocity in BEAS-2B cells. The more confined origin-centred trajectories and lower MSD profiles observed in GO:RGD treated U87 and U251 cells were qualitatively consistent with the mean-velocity results, whereas the trajectory and MSD patterns of BEAS-2B cells remained similar across treatment conditions.

#### Effect of GO:RGD on FAK Tyr3G7 phosphorylation

FAK Tyr397 phosphorylation was examined as an integrin-associated signalling endpoint after 24 h exposure to RGD (10 µg/mL), GO (50 µg/mL) or GO:RGD (50:10 µg/mL). Flow-cytometry analysis showed a significantly lower proportion of pFAK Tyr397-positive cells in GO:RGD-treated U87 cells than in untreated U87 cells (*p* < 0.05; Figure 7). Neither RGD nor GO alone produced a significant difference from the untreated control. No significant treatment-related differences were detected in U251 or BEAS-2B cells. The flow-cytometry gating strategy is shown in Figure S8. These findings indicate a cell-line-dependent association between GO:RGD treatment and a lower proportion of pFAK Tyr397-positive cells in U87 cells.

**Figure 7.**
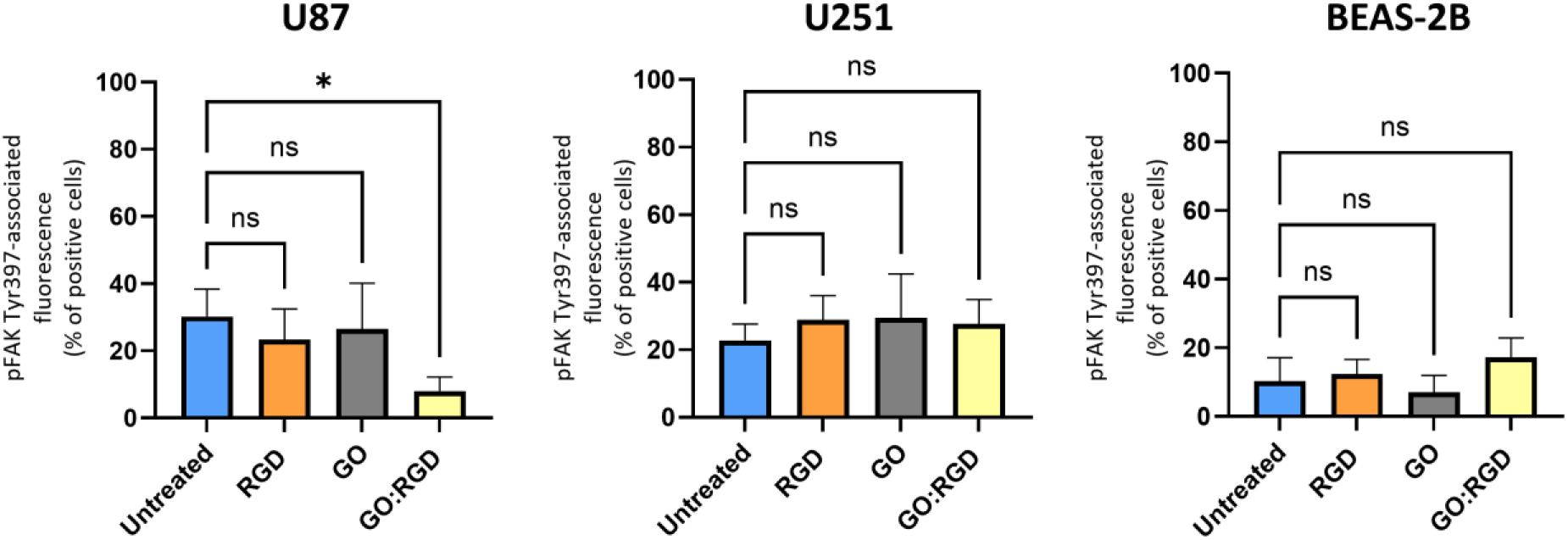
Effect of GO:RGD on FAK Tyr397 phosphorylation. U87, U251 and BEAS-2B cells were untreated or exposed to RGD (10 µg/mL), GO (50 µg/mL) or GO:RGD (50:10 µg/mL) for 24 h. FAK Tyr397 phosphorylation was assessed by flow cytometry. Data are presented as mean ± SD from three independent experiments (n = 3), with duplicate measurements averaged within each experiment. Statistical comparisons were performed separately for each cell line using one-way ANOVA followed by Dunnett’s multiple-comparisons test against the untreated control. The gating strategy is shown in Figure S8. *p < 0.05; ns, not significant.

## Discussion

This work explores the broader concept of using nanomaterial-cell membrane interactions as a platform for spatially controlling ligand presentation and modulating receptor-mediated signalling at the cell surface. As a proof-of-concept, we investigated whether the distinctive plasma-membrane association of GO in cancer cells could be used to position an RGD-containing ligand at the cell surface and modulate integrin-linked responses originating at the plasma membrane. A linear RGD-containing peptide was used as the model ligand, while cellular motility and downstream integrin-linked signalling were examined as functional readouts in glioblastoma cell lines. The physicochemical data supported non-covalent association of RGD with GO and retention of the sheet-like morphology and comparable size and thickness. In U87 and U251 glioblastoma cells, both GO and GO:RGD remained predominantly associated with the plasma membrane, whereas greater intracellular localisation was observed in BEAS-2B (non-cancer) cells. Functionally, GO:RGD reduced cell motility in two glioblastoma cell lines studied, but not in BEAS-2B cells, and a lower proportion of pFAK Tyr397-positive cells was detected in U87 cells. Together, these findings demonstrate cell-line-dependent functional consequence of membrane-localised GO, linking its differential plasma-membrane retention to reduction of glioblastoma cell motility and integrin-associated signalling.

Nanomaterial-mediated delivery to the plasma membrane remains an underexplored drug-delivery strategy. Carbon nanotube-, graphene- and DNA-based systems have been designed to present TRAIL to cell-surface death receptors [22–24]. However, these systems have focused principally on delivering an apoptosis-inducing cytokine. RGD-functionalised nanomaterials on the other hand generally exploit integrin recognition to promote receptor-mediated uptake and intracellular cargo delivery [18–20]. Here, GO:RGD retained the previously observed localisation behaviour of GO [9], remaining at the glioblastoma-cell periphery, positioning the non-covalently associated RGD ligand near the extracellular domains of integrins. This is particularly relevant because integrins recognise ligands extracellularly and undergo continuous internalisation and recycling [21]. Maintaining ligand availability at the cell surface rather than promoting its uptake may therefore provide a route to more sustained modulation of integrin-linked cellular processes. This approach also contrasts with our earlier defect-free graphene work, in which lysosomal internalisation was intentionally exploited for enzyme delivery [27]. Because membrane-associated GO can itself alter plasma-membrane tension and downstream signalling [28], it also represents an active nano-bio interface.

Because the plasma membrane-associated behaviour exploited in this study depends on the physicochemical properties of GO [9], it was essential to first determine whether RGD association preserved the key characteristics of the starting GO nanosheets, especially size and thickness. The physicochemical characterisation supports the formation of a dynamic GO:RGD assembly. Increased nitrogen content detected by XPS and the lower-angle XRD peak following purification indicate peptide association with GO, while AFM confirmed retention of the sheet-like morphology. The decrease in GO-associated peptide over 24 h in water supports a dynamic non-covalent GO:RGD assembly, distinguishing it from covalently functionalised GO:RGD constructs [29,30]. While covalent attachment can enhance ligand stability, it may alter GO surface properties, whereas non-covalent association may preserve greater conformational flexibility. DLS measurements should be interpreted cautiously due to the limitations of modelling plate-like structures, but they provide comparative insight into dispersion behaviour. Overall, peptide association was achieved without substantially altering nanosheet morphology.

The observed cellular effects arise from the combined influence of peptide presentation and GO-cell interactions. Association with GO may increase the effective local concentration of RGD at the plasma membrane or present multiple ligand molecules within a confined surface region. Nanoparticle-based presentation can enhance ligand-receptor interactions through increased local density [19]. In addition, GO itself may influence plasma-membrane properties [28] and cytoskeletal dynamics [31–33], thereby modifying how cells respond to the associated peptide. GO:RGD should therefore be considered a formulation-dependent system in which biological activity reflects both peptide-mediated and material-driven effects.

The effects on cell movement provide functional insight into the combined influence of GO-cell interactions and RGD presentation. Mean cell velocity quantifies how rapidly individual cells move, whereas trajectory and mean-square-displacement analyses describe how far and how freely they travel over time. The more confined trajectories and lower displacement profiles therefore qualitatively support the reduction in velocity, together indicating restricted cell movement. In U87 cells, GO:RGD reduced mean velocity whereas GO and free RGD did not, and this effect was accompanied by reduced pFAK-associated fluorescence. This pattern is consistent with an RGD-dependent contribution to integrin-linked signalling that becomes evident when the peptide is presented in association with membrane-localised GO. In contrast, both GO and GO:RGD reduced mean velocity in U251 cells, without a corresponding change in pFAK. This suggests a stronger GO-dependent component in U251 cells, potentially involving cytoskeletal or metabolic pathways not captured by the pFAK endpoint. Previous studies have shown that GO can impair cell migration through effects on actin organisation and mitochondrial function [31–33], supporting this interpretation.

These differences also indicate that the proportion of integrin-positive cells alone does not determine sensitivity to the formulation. Although U87 cells exhibited higher fractions of αvβ3-, αvβ5- and α5β1-positive cells, U251 cells still responded in the motility assay despite lower integrin-positive fractions. Furthermore, the linear RGD sequence can interact with multiple integrin subtypes and is not selective for αvβ3 [16,17,20]. While the presence of GO and αvβ3 staining within the same peripheral regions supports proximity at the cell surface, it does not identify αvβ3 as the sole mediator of the observed effects. Variations in integrin repertoire, adhesion dynamics, cytoskeletal organisation and GO–membrane interactions likely contribute to the cell-specific responses.

The behaviour of BEAS-2B cells further highlights the importance of material localisation. In these cells, GO and GO:RGD showed greater intracellular localisation and did not produce significant changes in motility or pFAK-associated fluorescence. This is explained by GO:RGD internalisation reducing the persistence of the formulation within the extracellular receptor environment. Further studies using non-malignant neural cells, as well as patient-derived glioblastoma models are needed to strengthen this conclusion.

The pFAK data provide further mechanistic insight. Phosphorylation of FAK at Tyr397 integrates signals from integrins as well as growth-factor and cytokine receptors [25,34–37]. The reduced pFAK-associated signal observed in U87 cells following GO:RGD treatment may reflect altered integrin-mediated adhesion, secondary effects of reduced motility, or both. The absence of a pFAK response in U251 cells despite reduced velocity indicates that FAK signalling is not the sole pathway involved. Alternative mechanisms, including Rho-family GTPase signalling, integrin recycling and actomyosin regulation, warrant further investigation [21,38]. Time-resolved studies would also be valuable, as a single 24 h endpoint may not capture transient signalling dynamics.

Several limitations should be considered. The study did not directly assess peptide accessibility, receptor occupancy, and quantitative co-localisation. The study also used two established glioblastoma cell lines and one non-cancerous comparator of a different tissue origin. Mean cell velocity served as the primary motility endpoint, supported by trajectory and MSD analyses, but these two-dimensional measurements do not directly reflect tumour invasion or metastatic potential. Future work incorporating scrambled or non-binding control peptides and three-dimensional glioblastoma models will be important to further define mechanism and biological relevance.

This study provides a proof-of-concept that the cell-line-dependent localisation of GO can be exploited to present a surface-active ligand within the plasma membrane environment. The distinct responses observed in U87 and U251 cells indicate that GO:RGD activity likely reflects an interplay between peptide presentation, integrin biology and the intrinsic effects of GO. This framework offers a basis for developing membrane-associated nanomaterial platforms for modulating cell-surface receptors, while highlighting the need for further mechanistic and translational investigations.

## Conclusion

This study investigated whether graphene oxide can function as a plasma membrane-localised nanoplatform for the presentation of bioactive ligands and the modulation of receptor-associated signalling in cancer cells. Integrins were used as the model receptor system, the RGD motif as a functional ligand known to interfere with integrin activity, glioblastoma cell lines as the disease-relevant model, and cell motility together with downstream signalling as primary readouts.

Non-covalent association of GO with RGD preserved the key physicochemical properties of the nanosheets, namely size and thickness. GO retained a predominantly plasma membrane-associated localisation in both U87 and U251 glioblastoma cells, while being efficiently internalised by non-cancerous BEAS-2B cell line. GO:RGD exposure reduced cell motility in both glioblastoma lines and reduced the proportion of pFAK Tyr397-positive cells in U87, whereas no significant effects were observed in BEAS-2B cells. The differing responses between U87 and U251 cells suggest that GO:RGD-associated effects arise from a combination of ligand presentation, integrin signalling context, and intrinsic cell-line-specific interactions with GO.

Collectively, these findings reposition graphene oxide from an intracellular nanocarrier to an active, membrane-associated nano–bio interface capable of influencing cell-surface receptor function. Further work is required to directly confirm ligand accessibility and receptor engagement in more physiologically relevant glioblastoma cellular models and *in vivo*. This work opens the way to the rational design of graphene-based platforms for spatially controlled modulation of plasma membrane receptor signalling in cancer cells.

## Methods

### Non-covalent complexation of GO with RGD

Endotoxin-free GO was produced in-house and fully characterised, as previously described [4]. For GO:RGD complex preparation, the RGD peptide (GRGDS; catalogue 4008998.0025; Bachem, supplied by Cambridge Bioscience, UK) was added to GO nanosheets in ultrapure water, corresponding to a 10:2 GO:RGD mass ratio. The mixture was stirred at 500 rpm for 30 min at room temperature. For physicochemical characterisation, the GO:RGD complex was purified to remove unbound peptide prior to analysis, using centrifugal filtration (2,000 × g for 5 min; Amicon Ultra-4 Centrifugal Filter Unit, 100 kDa, 4 mL; EMD Millipore, UK). Four sequential centrifugal-filtration washes were performed, with the retained GO:RGD restored to the initial volume with water after each wash, to ensure efficient removal of the unbound peptide before characterisation.

Unbound RGD was quantified using a TNBSA (2,4,6-trinitrobenzene sulfonic acid) assay (TS-28997; Thermo Fisher Scientific, UK). Separate GO:RGD preparations were maintained in water for 0, 4 or 24 h following complexation. At each time point, samples underwent four sequential centrifugal-filtration washes, and the combined filtrates were analysed by TNBSA to determine unbound RGD. The filtrate from each wash was restored to the initial volume with water. The calibration curve was prepared using RGD standards prepared in 0.1 M sodium bicarbonate solution (pH 8.5; S5761-1KG, Sigma-Aldrich, UK) at concentrations ranging from 0.5 to 50 µg/mL. A 0.01% (w/v) TNBSA solution (250 μL) was added to each sample. Samples were mixed at 300 rpm for 2 h at 37 °C. The reaction was terminated by the addition of 250 μL of 10% (w/v) sodium dodecyl sulfate (SDS; L5750-100G, Sigma-Aldrich, UK), followed by 125 μL of 1 M hydrochloric acid (37%; 258148, Sigma-Aldrich, UK). Samples were aliquoted into clear 96-well plates (Greiner Bio-One, UK), and absorbance was measured at 340 nm using an Omega microplate reader. Background absorbance was subtracted from the sample absorbance. Negative background-corrected readings were set to zero. The absorbance-concentration relationship was plotted in OriginPro 9.1 to generate the calibration curve. The unbound RGD concentration was determined from the TNBSA absorbance of the filtrates using the slope of the calibration curve. The initial amount of RGD added was defined as 100%, and the GO-associated fraction was calculated as:

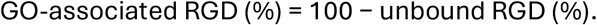

Three independently prepared samples were analysed, with absorbance measured in technical triplicate. The four sequential wash fractions were treated as fractions from the same preparation rather than as independent replicates.

### Dynamic light scattering (DLS)

Hydrodynamic size and polydispersity index (PDI) were measured at 0, 4 and 24 h using a Zetasizer Nano ZS (Malvern Instruments, UK). One millilitre of each sample (20 μg/mL) was transferred to a folded capillary zeta cell (DTS1070; Malvern Instruments, UK). Three independently prepared samples were analysed at room temperature, with each sample measured in technical triplicate. Data are reported as mean ± standard deviation (SD). All samples were monitored visually by capturing images at every time point.

### Zeta-potential measurements

Electrophoretic mobility was measured at 0, 4 and 24 h using a Zetasizer Nano ZS (Malvern Instruments, UK). One millilitre of GO or GO:RGD (20 μg/mL) was transferred to a folded capillary zeta cell (DTS1070; Malvern Instruments, UK). Three independently prepared samples were analysed at room temperature, with each sample measured in technical triplicate. Data are reported as mean ± SD.

### pH measurements

The pH of all samples (1 mL) was measured using a FiveEasy FE20 benchtop pH meter equipped with an InLab Micro Pro pH electrode (30224732, METTLER TOLEDO Ltd., Leicester, UK). Prior to conducting the measurement, the instrument was calibrated using standard buffer solutions (pH 4.00, 7.00, and 10.00) at room temperature. The electrode was rinsed with deionised water between measurements, and the pH values were recorded after the readings had stabilised.

### X-ray photoelectron spectroscopy (XPS)

Twenty microlitres of each sample was drop-cast onto a 5 × 5 mm Si wafer (Ted Pella) to form a thin film. Spectra were acquired using a Phoibos 150 electron spectrometer (SPECS GmbH) coupled to a hemispherical analyser under ultrahigh-vacuum conditions, with an Al Kα X-ray source (hν = 1486.74 eV). Measurements were performed at the ICN2 Photoemission Spectroscopy Facility. Charge effects were corrected by referencing the adventitious-carbon C 1s peak to 284.6 eV. Deconvolution of the C 1s, O 1s and N 1s spectra was performed after Shirley background subtraction using Gaussian–Lorentzian (70:30) peak fitting, with the full width at half maximum constrained between 0.5 and 2 eV. CasaXPS software was used for data analysis.

### X-ray diffraction (XRD)

Two hundred microlitres of each sample was drop-cast onto a Si holder and left to dry in an oven at 50 °C. Spectra were acquired using a Malvern PANalytical X’Pert Pro MPD diffractometer over a 2θ scan range of 5°–60°. Measurements were performed at the ICN2 XRD Facility using a ceramic X-ray tube with a Cu Kα anode (λ = 1.540 Å) and an X’Celerator solid-state detector. Spectra were analysed using X’Pert HighScore (version 2.2c [2.2.3]) and plotted in OriginPro 9.1.

### Atomic force microscopy (AFM)

GO and GO:RGD were characterised using a MultiMode 8-HR atomic force microscope (Bruker, UK) in ScanAsyst® mode. Measurement was performed at the Bio-AFM Facility of the University of Manchester. Silicon-coated cantilevers (Bruker, UK) with a resonance frequency of 60 kHz and a force constant of 0.5 N m⁻¹ were used to acquire AFM height images. For sample preparation, cleaved mica was mounted on a 12 mm iron disc using carbon tape, and 20 µL of 0.01% poly-L-lysine (Sigma-Aldrich, UK) was drop-cast onto the mica. After 1 min, the mica was washed with 1 mL water, followed by drop-casting 20 μL GO:RGD (50 μg/mL). The same procedure was used for GO. Unbound material was removed by two washes with 1 mL water. The samples were left to dry overnight, and the acquired images were processed using Gwyddion 7.1.

### Cell culture

Immortalised human bronchial epithelial BEAS-2B cells (CRL-9609, ATCC, LGC standards, UK) and human U251 glioblastoma cells (kindly provided by Prof. Karen Kirkby, Precise Group, NHS Proton Therapy Centre at The Christie) were maintained in RPMI-1640 medium. Human U87 MG glioblastoma cells (HTB-14; ATCC, UK) were maintained in DMEM (Sigma-Aldrich, UK). To obtain complete cell culture medium, the RPMI-1640 or DMEM was supplemented with 10% FBS (Gibco, UK), 100 U/mL and 100 µg/mL penicillin and streptomycin, respectively. Cells were maintained at 37 °C in a humidified 5% CO₂ incubator and passaged at approximately 80% confluence using 0.05% trypsin-EDTA (Sigma-Aldrich, UK). Trypsin activity was quenched with 10% FBS.

### Cell culture treatments

For biological experiments, GO:RGD preparation was diluted directly into complete culture medium. All GO:RGD preparations used a fixed 10:2 GO:RGD mass ratio; total amounts and volumes were scaled according to experimental requirements. A free-RGD treatment condition always matched the starting peptide concentration used for the complexation preparation. Cells were seeded 24 h before treatment in Cellview™ four-well dishes for confocal and time-lapse experiments or 12-well Corning™ plates for flow-cytometry experiments. Cells were treated at 60%–80% confluence in the complete cell culture medium. Cells were maintained at 37 °C in a humidified 5% CO₂ incubator.

### Flow cytometry

#### Annexin V/propidium iodide assay

Cells were treated with RGD (0, 10, 25, 50, 75 or 100 μg/mL; 1 mL per well) for 24 h in the complete growth medium. After 24 h the cell media was collected and cells were detached using 0.05% trypsin-EDTA (300 μL per well, 10 min), followed by neutralisation using 10% FBS (30 μL per well), and collected in 1.5 mL microcentrifuge tubes. Cells were centrifuged at 220 × g for 5 min and resuspended in 1× Annexin-binding buffer (V13246, Thermo Fisher, UK; 200 μL per tube). Cells were stained with Annexin V (A13201, Thermo Fisher, UK; 1 μL per tube) for 20 min. Samples were kept on ice until analysis using a FACSVerse flow cytometer (BD Biosciences). Propidium iodide (P4864, Sigma-Aldrich, UK; 1 μL per tube) was added immediately before acquisition. Annexin V was detected in the FITC-A channel using 488 nm excitation and a 530/30 nm band-pass filter; propidium iodide was detected in the PE-A channel using 488 nm excitation and a 574/26 nm band-pass filter.

#### Integrin expression analysis

All cell lines were seeded in 12-well plates and detached with 0.05% trypsin-EDTA (300 μL per well, 10 min), followed by neutralisation with 10% FBS (30 μL per well), and collected in 1.5 mL microcentrifuge tubes. Cells were centrifuged at 220 × g for 5 min and resuspended in 200 μL of 2% paraformaldehyde (PFA; Thermo Fisher Scientific, UK) for 30 min at room temperature. After fixation, 500 μL FACS buffer (2 mM EDTA, Invitrogen, UK, and 0.5% BSA, Gibco, UK, in calcium- and magnesium-free PBS, Sigma-Aldrich, UK) was added to each sample, and cells were centrifuged at 220 × g for 5 min. Cells were resuspended in FACS buffer and stained with mouse monoclonal anti-αvβ3 integrin antibody (clone LM609; ab190147, Abcam, UK) at 1:500 for 1 h on ice. Cells were washed twice with FACS buffer and stained with goat anti-mouse Alexa Fluor™ 488 secondary antibody (A-11001, Invitrogen, UK) at 1:500 for 30 min on ice. Cells were washed twice, resuspended in fresh FACS buffer and stored on ice until analysis using a FACSVerse flow cytometer. Alexa Fluor 488 was detected using 488 nm excitation and a 530/30 nm band-pass filter. The same procedure was followed for αvβ5 (mouse monoclonal clone P1F6; ab177004, Abcam, UK) and α5β1 (recombinant chimeric rabbit clone M200 [volociximab]; ab275977, Abcam, UK). Goat anti-mouse Alexa Fluor™ 488 was used for αvβ5, whereas goat anti-rabbit Alexa Fluor™ 488 (A-11008, Invitrogen, UK) was used for α5β1.

#### FAK Tyr3G7 phosphorylation analysis

All cell lines were seeded in 12-well plates and exposed to medium alone (untreated), RGD (10 μg/mL), GO (50 μg/mL) or GO:RGD (50:10 μg/mL); 1 mL per well for 24 h in the appropriate complete growth medium. Cells were detached using 0.05% trypsin-EDTA (300 μL per well, 10 min), neutralised using 10% FBS (30 μL per well), collected in 1.5 mL microcentrifuge tubes and centrifuged at 220 × g for 5 min. Cells were resuspended in 2% PFA (200 μL per tube) for 30 min at room temperature. After fixation, cells were washed with PBS and permeabilized with 0.1% Triton X-100 in PBS for 15 min at room temperature. Cells were then washed with 500 μL of FACS buffer and centrifuged at 220 × *g* for 5 min. They were then resuspended in FACS buffer and stained with anti-phospho-FAK (Tyr397) primary antibody (rabbit; 44-624G, Thermo Fisher Scientific, Invitrogen, UK) at 1:500 for 1 h on ice. Cells were washed twice and stained with goat anti-rabbit Alexa Fluor™ 488 secondary antibody (A-11008, Invitrogen, UK) at 1:500 for 30 min on ice. Cells were washed twice, resuspended in fresh FACS buffer and stored on ice until analysis using a FACSVerse flow cytometer. Alexa Fluor 488 was detected using 488 nm excitation and a 530/30 nm band-pass filter.

#### Optical microscopy

Optical microscopy images were acquired using a Zeiss Primovert inverted microscope in transmitted-light mode. Samples were imaged in bright field using 10× and 20× objectives. Images were analysed using ZEN imaging software.

### Confocal microscopy

#### Interactions with the plasma membrane/uptake analysis

Cells were treated with complete cell culture medium alone (untreated), RGD (5 μg/mL), GO (25 μg/mL) or GO:RGD (25:5 μg/mL) in 0.5 mL final volume per well for 24 h. At 24 h post-treatment, supernatants were removed and cells were stained with CellMask™ Green plasma-membrane stain (C37608, Thermo Fisher Scientific, UK), prepared in complete medium at 1:2,500. Live-cell imaging was performed using a Zeiss 780 confocal laser-scanning microscope with a 40× oil-immersion objective. Images were processed using ZEN software. CellMask Green was detected using 488/520 nm excitation/emission, while GO was visualised through its intrinsic fluorescence using 594 nm excitation and a 620–690 nm emission window.

#### Interaction of GO:RGD with αvβ3 integrin

Cells were treated with complete cell culture medium alone (untreated), RGD (5 μg/mL), GO (25 μg/mL) or GO:RGD (25:5 μg/mL) in 0.5 mL final volume per well for 24 h. Supernatants were removed, and cells were washed with pre-warmed PBS +/+ (0.5 mL per well; Sigma-Aldrich, UK) and fixed with 4% PFA (0.5 mL per well) for 15 min. Cells were blocked with 5% goat serum (Sigma-Aldrich, UK) in PBS for 1 h at room temperature and incubated with mouse monoclonal anti-αvβ3 integrin antibody (clone LM609; ab190147, Abcam, UK) at 1:500 overnight at 4 °C. The following day, cells were washed three times with PBS, with the final wash extended to 10 min, and stained with goat anti-mouse Alexa Fluor™ 488 secondary antibody (A-11001, Invitrogen, UK) at 1:500 for 2 h at room temperature in the dark. Cells were washed three times, with the final wash extended to 10 min, and 0.5 mL fresh PBS was added to each well. Cells were imaged using a Zeiss 780 confocal laser-scanning microscope with a 40× objective. Images were processed using ZEN software. Alexa Fluor 488 was detected at 488/520 nm, while GO was detected using 594 nm excitation and a 620–690 nm emission window.

#### Time-lapse microscopy: video acquisition

Cells were stained with Hoechst 33342 (1 μg/mL; 0.5 mL per well; 10 min; 62249, Thermo Fisher Scientific, UK). Cells were then washed once with corresponding complete medium and exposed to complete medium (untreated), RGD (5 μg/mL), GO (25 μg/mL) or GO:RGD (25:5 μg/mL) in 0.5 mL per well. Cells were imaged continuously for 24 h, with one frame acquired every 11 min, using a Zeiss 710 confocal laser-scanning microscope in tile mode. Acquired videos were analysed using ZEN, CellTracker and DiPer software, as described in [9]. Individual cells were tracked manually using CellTracker version 1.1. Live-cell time-lapse videos were exported using ZEN Black software (Carl Zeiss Microscopy) and loaded into CellTracker. The 16-tile video from each condition was cropped to four tiles, and cells were tracked manually across all 135 frames. Each cell was tracked by clicking on its nucleus in every frame. Twenty-five cells per condition were tracked in each of three independent experiments, giving 75 tracked cells per condition in total. Average velocity (µm/min) and xy coordinates were calculated using the Statistics tab in CellTracker after entering a spatial calibration of 0.346 µm per pixel and a temporal interval of 11 min per frame. The xy-coordinate data were imported into DiPer Excel macro-enabled workbooks to generate origin-centred trajectory plots and mean-square-displacement (MSD) curves. Velocity data were plotted using GraphPad Prism 9.0.1.

#### Statistical analysis

All experiments were repeated at least two times with duplicates or triplicates for each condition. Statistical analyses were performed using GraphPad Prism 9.0.1. Data are presented as mean ± SD or SEM, as indicated. Comparisons among untreated, RGD, GO and GO:RGD groups were performed separately within each cell line using one-way ANOVA followed by Dunnett’s multiple-comparisons test against the untreated control. For motility analysis, the mean velocity of 25 tracked cells within each independent experiment represented one experimental value. Technical replicates from flow-cytometry experiments were averaged to produce one value per independent experiment. Differences were considered statistically significant at p < 0.05. *p < 0.05, **p < 0.01, ***p < 0.001 and ****p < 0.0001; ns, not significant.

## Author Contributions

S.V. conceived and supervised the work. A.K. and S.V. designed all the experiments. A.K. performed the experiments, analysed the data, and wrote the manuscript. N.L. provided the materials and supervised the preparation of GO complexes with RGD. K.K. supervised preparation and characterisation of GO. The manuscript was edited by A.K., S.V. and N.L. with the input from K.K.

## Acknowledgements

A.K. would like to acknowledge the studentship from the Engineering and Physical Sciences Research Council (EPSRC) Centre for Doctoral Training programme (Graphene NOWNANO CDT). The authors would like to acknowledge Dr. Luis M. Arellano and Ms Irene Rebollido Pedrido for their contribution to the synthesis and characterisation of the specific GO batch used in this study. The authors would like to acknowledge Ms Hafsah Shah who assisted with the performance and analysis of the time-lapse motility experiments, Ms Ariadna Fuertes Gassio who carried out the cytotoxicity experiments and Dr. Tommaso Battisti for acquiring the XPS and XRD data. The ICN2 has been supported by the Severo Ochoa Centres of Excellence programme [SEV-2017-0706] and is currently supported by the Severo Ochoa Centres of Excellence programme, Grant CEX2021-001214-S, both funded by MCIN and MCIU/AEI/10.13039.501100011033. The authors would like to acknowledge the staff of the Manchester Bioimaging Facility, and specifically Dr. David Spiller for his assistance and guidance regarding the confocal and time-lapse video microscopy performed in this study. The authors also wish to thank Dr. Nigel Hodson from the Bio-AFM Facility for assistance and advice regarding the AFM instrumentation. The authors would like to acknowledge the flow cytometry Core Facility of the University of Manchester and specifically, Dr. Gareth Howell for his assistance regarding the flow cytometry studies conducted in this study. Illustrations were created with BioRender.com.

## References

[1] Muzyka, R., Kwoka, M., Smędowski, Ł., Díez, N. & Gryglewicz, G. Oxidation of graphite by different modified Hummers methods. New Carbon Materials 32, 15–20 (2017).

[2] Mao, H. Y. et al. Graphene: Promises, facts, opportunities, and challenges in nanomedicine. Chemical Reviews 113, 3407–3424 (2013).

3. Georgakilas, V. Functionalization of Graphene. Wiley-VCH (2014).

[4] Rodrigues, A. F. et al. A blueprint for the synthesis and characterisation of thin graphene oxide with controlled lateral dimensions for biomedicine. 2D Materials 5, 035020 (2018).

[5] Liu, J., Cui, L. & Losic, D. Graphene and graphene oxide as new nanocarriers for drug delivery applications. Acta Biomaterialia 9, 9243–9257 (2013).

[6] Itoo, A. M. et al. Multifunctional graphene oxide nanoparticles for drug delivery in cancer. Journal of Controlled Release 350, 26–59 (2022).

[7] Zhang, B., Wei, P., Zhou, Z. & Wei, T. Interactions of graphene with mammalian cells: Molecular mechanisms and biomedical insights. Advanced Drug Delivery Reviews 105, 145–162 (2016).

[8] Chen, Y. et al. Dynamic interactions and intracellular fate of label-free, thin graphene oxide sheets within mammalian cells: Role of lateral sheet size. Nanoscale Advances 3, 4166–4185 (2021).

[9] Chen, Y., Rosano, V., Lozano, N. et al. Interplay between material properties and cellular effects drives distinct pattern of interaction of graphene oxide with cancer and non-cancer cells. Journal of Nanobiotechnology 23, 393 (2025).

[10] Hynes, R. O. Integrins: Bidirectional, allosteric signaling machines. Cell 110, 673–687 (2002).

[11] Hamidi, H. & Ivaska, J. Every step of the way: Integrins in cancer progression and metastasis. Nature Reviews Cancer 18, 533–548 (2018).

[12] Echavidre, W., Picco, V., Faraggi, M. & Montemagno, C. Integrin-αvβ3 as a therapeutic target in glioblastoma: Back to the future? Pharmaceutics 14, 1053 (2022).

[13] Bello, L. et al. Alpha(v)beta3 and alpha(v)beta5 integrin expression in glioma periphery. Neurosurgery 49, 380–390 (2001).

[14] Janouskova, H. et al. Integrin α5β1 plays a critical role in resistance to temozolomide by interfering with the p53 pathway in high-grade glioma. Cancer Research 72, 3463–3470 (2012).

[15] Desgrosellier, J. S. & Cheresh, D. A. Integrins in cancer: Biological implications and therapeutic opportunities. Nature Reviews Cancer 10, 9–22 (2010).

[16] Alipour, M. et al. Recent progress in biomedical applications of RGD-based ligand: From precise cancer theranostics to biomaterial engineering: A systematic review. Journal of Biomedical Materials Research Part A 108, 839–850 (2020).

[17] Roxin, Á. & Zheng, G. Flexible or fixed: A comparative review of linear and cyclic cancer-targeting peptides. Future Medicinal Chemistry 4, 1601–1618 (2012).

[18] Yang, X., et al. cRGD-functionalized, DOX-conjugated, and 64Cu-labeled superparamagnetic iron oxide nanoparticles for targeted anticancer drug delivery and PET/MR imaging. Biomaterials 32, 4151–4160 (2011).

[19] Montet, X., Funovics, M., Montet-Abou, K., Weissleder, R. & Josephson, L. Multivalent effects of RGD peptides obtained by nanoparticle display. Journal of Medicinal Chemistry 49, 6087–6093 (2006).

[20] Rios De La Rosa, J. M., et al. Microfluidic-assisted preparation of RGD-decorated nanoparticles: Exploring integrin-facilitated uptake in cancer cell lines. Scientific Reports 10, 14505 (2020).

[21] White, D. P., Caswell, P. T. & Norman, J. C. αvβ3 and α5β1 integrin recycling pathways dictate downstream Rho kinase signaling to regulate persistent cell migration. Journal of Cell Biology 177, 515–525 (2007).

[22] Zakaria, A. B. et al. Nanovectorization of TRAIL with single wall carbon nanotubes enhances tumor cell killing. Nano Letters 15, 891–895 (2015).

[23] Jiang, T. et al. Furin-mediated sequential delivery of anticancer cytokine and small-molecule drug shuttled by graphene. Advanced Materials 27, 1021–1028 (2015).

[24] Sun, W. et al. Transformable DNA nanocarriers for plasma membrane targeted delivery of cytokine. Biomaterials 96, 1–10 (2016).

[25] Schaller, M. D. Cellular functions of FAK kinases: Insight into molecular mechanisms and novel functions. Journal of Cell Science 123, 1007–1013 (2010).

[26] Vranic, S. et al. Live imaging of label-free graphene oxide reveals critical factors causing oxidative-stress-mediated cellular responses. ACS Nano 12, 1373–1389 (2018).

[27] Chen, Y. et al. Defect-free graphene enhances enzyme delivery to fibroblasts derived from patients with lysosomal storage disorders. Nanoscale 15, 9348–9364 (2023).

[28] Ogene, L. et al. Graphene oxide activates canonical TGFβ signalling in a human chondrocyte cell line via increased plasma membrane tension. Nanoscale 16, 5653–5664 (2024).

[29] Jagiełło, J. et al. Adhesive properties of graphene oxide and its modification with RGD peptide towards L929 cells. Materials Today Communications 26, 102056 (2021).

[30] Eckhart, K. E., Holt, B. D., Laurencin, M. G. & Sydlik, S. A. Covalent conjugation of bioactive peptides to graphene oxide for biomedical applications. Biomaterials Science 7, 3876–3885 (2019).

[31] Zhou, H. et al. The inhibition of migration and invasion of cancer cells by graphene via the impairment of mitochondrial respiration. Biomaterials 35, 1597–1607 (2014).

[32] Yu, Q., Zhang, B., Li, J. & Li, M. The design of peptide-grafted graphene oxide targeting the actin cytoskeleton for efficient cancer therapy. Chemical Communications 53, 11433–11436 (2017).

[33] Tian, X. et al. Graphene oxide nanosheets retard cellular migration via disruption of actin cytoskeleton. Small 13, 1602133 (2017).

[34] Kleinschmidt, E. G. & Schlaepfer, D. D. Focal adhesion kinase signaling in unexpected places. Current Opinion in Cell Biology 45, 24–30 (2017).

[35] Tan, X. et al. Focal adhesion kinase: from biological functions to therapeutic strategies. Experimental Hematology & Oncology 12, 83 (2023).

[36] Sieg, D. J. et al. FAK integrates growth-factor and integrin signals to promote cell migration. Nature Cell Biology 2, 249–256 (2000).

[37] Nuñez, R. E., del Valle, M. M., Ortiz, K., Almodovar, L. & Kucheryavykh, L. Microglial cytokines induce invasiveness and proliferation of human glioblastoma through Pyk2 and FAK activation. Cancers 13, 6160 (2021).

[38] Cox, E. A., Sastry, S. K. & Huttenlocher, A. Integrin-mediated adhesion regulates cell polarity and membrane protrusion through the Rho family of GTPases. Molecular Biology of the Cell 12, 265–277 (2001).

