## Supporting Information for "Plasma membrane-associated graphene oxide as a platform for modulating signalling through cell-surface receptors: an integrin-focused proof-of-concept study"

| Physicochemical properties | Technique | GO |
|---|---|---|
| Lateral dimension | AFM | 25 nm–1.5 μm (95% < 475 nm) |
| | SEM | 50 nm–1.9 μm (95% < 850 nm) |
| Thickness | AFM | 1–2 nm |
| Optical properties | Absorbance | $A_{230}$ = 0.053 × C_GO (μg/mL) |
| Degree of defects (I_D/I_G) | Raman spectroscopy | 1.14 ± 0.03 |
| Surface charge | ζ-Potential | -52.1 ± 0.4 mV |
| Chemical Composition (Purity) | XPS | C: 72.2%, O: 25.0%, S: 1.2%, B: 1.6% (97.2%) |

**Table S1. Physicochemical properties of the starting GO nanosheets dispersed in water.** Lateral dimensions were determined by atomic force microscopy (AFM) and scanning electron microscopy (SEM), nanosheet thickness by AFM, optical properties by absorbance spectroscopy, structural defects by Raman spectroscopy, surface charge by zeta-potential measurement and elemental composition by X-ray photoelectron spectroscopy (XPS). Values are presented as mean ± SD where applicable.

| Time | Condition | Replicate 1 | Replicate 2 | Replicate 3 | Mean Abs | SD.S (+/-) | Peptide (µg/mL) | Percentage % of free peptide | Percentage of complexation (%) |
|---|---|---|---|---|---|---|---|---|---|
| 0 h | GO | 0,0005 | 0 | 0 | 0,000167 | 0,000289 | 0,053419 | 0,244698206 | NA |
| | RGD | 0,059 | 0,06 | 0,081 | 0,068111 | 0,011403 | 21,83048 | 100 | 0 |
| | GO:RGD | 0,020 | 0,02 | 0,02457 | 0,020612 | 0,003797 | 6,606481 | 30,26264274 | 69,74 |
| 4h | GO | 0,00033 | 0 | 0,005 | 0,002332 | 0,002405 | 0,747507 | 3,144569293 | NA |
| | RGD | 0,073 | 0,06 | 0,086 | 0,074167 | 0,011439 | 23,77137 | 100 | 0 |
| | GO:RGD | 0,035 | 0,01 | 0,026733333 | 0,024133 | 0,008955 | 7,735007 | 32,53917608 | 67,5 |
| 24 h | GO | 0,00006 | 0 | 0,0055 | 0,001853 | 0,003158 | 0,594017 | 2,523831139 | NA |
| | RGD | 0,073 | 0,06 | 0,086 | 0,073433 | 0,012356 | 23,53632 | 100 | 0 |
| | GO:RGD | 0,038 | 0,03 | 0,043 | 0,037633 | 0,00535 | 12,06197 | 51,24829778 | 48,8 |

**Table S2. Quantification of unbound and GO-associated RGD using the TNBSA assay.** GO-only, free-RGD and GO:RGD samples prepared at a fixed 10:2 GO mass ratio were analysed at 0, 4 and 24 h. Background-corrected absorbance values from three independently prepared samples were averaged and converted to RGD concentrations using the calibration curve shown in Figure S2. At each time point, unbound RGD was expressed relative to the corresponding free-RGD control, defined as 100%, and the GO-associated fraction was calculated as 100 − unbound RGD (%). Data are presented as mean ± SD. NA, not applicable.

| Time | Condition | Wash 1 | Wash 2 | Wash 3 | Wash 4 | Total Abs | Calculated peptide (µg/mL) Abs/Slope | Percentage % of free peptide |
|---|---|---|---|---|---|---|---|---|
| 0 h | GO | 0 | 0 | 0 | 0 | 0 | 0 | NA |
| | RGD | 0,064 | 0 | 0 | 0 | 0,064 | 20,51282 | 100 |
| | GO:RGD | 0,011 | 0,006 | 0 | 0 | 0,017 | 5,448718 | 26,6 |
| 4h | GO | 0 | 0 | 0 | 0,001667 | 0,0017 | 0,534188 | NA |
| | RGD | 0,0536 | 0,005 | 0,0045 | 0 | 0,0632 | 20,24573 | 100 |
| | GO:RGD | 0,0073 | 0,0015 | 0 | 0,005333 | 0,0142 | 4,540491 | 22,4 |
| 24 h | GO | 0 | 0 | 0 | 0 | 0 | 0 | NA |
| | RGD | 0,057 | 0 | 0,004 | 0,0003 | 0,0613 | 19,64744 | 100 |
| | GO:RGD | 0,007 | 0,006 | 0,001 | 0,0183 | 0,0323 | 10,35256 | 52,7 |

**Table S3. Recovery of unbound RGD during sequential centrifugal-filtration washes.** TNBSA absorbance values are shown for the four sequential filtrate fractions collected from GO-only, free-RGD and GO:RGD preparations at 0, 4 and 24 h. Total absorbance was calculated by summing the four wash fractions and converted to RGD concentration using the calibration-curve slope. Unbound RGD was expressed relative to the corresponding free-RGD control. The four wash fractions originated from the same preparation and were not treated as independent replicates. NA, not applicable.

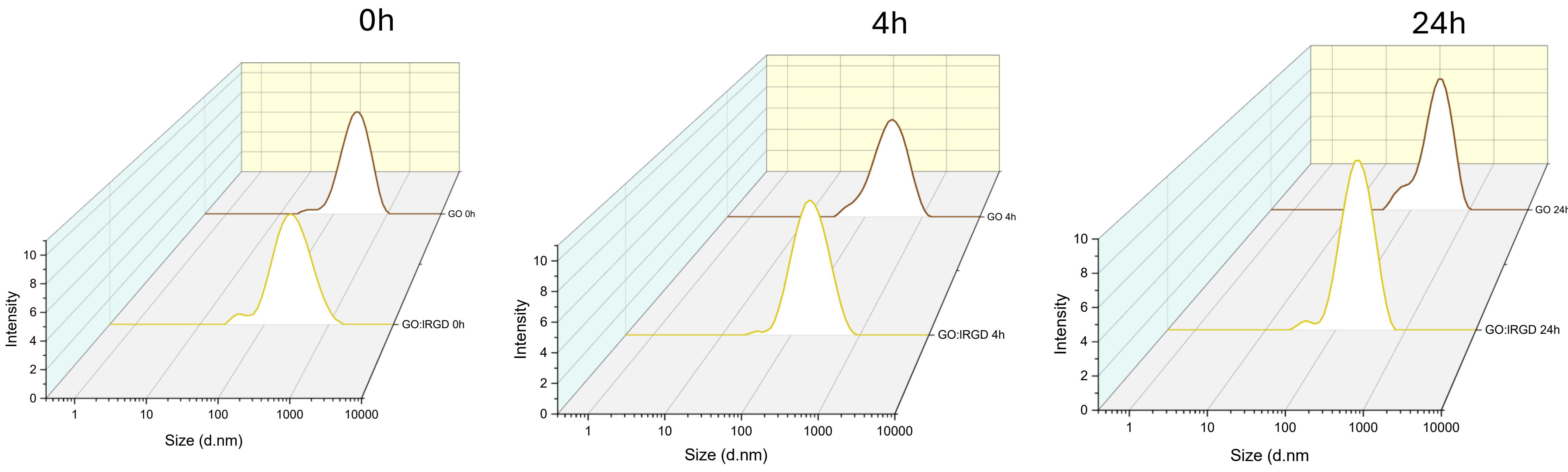


**Figure S1. Intensity-weighted apparent hydrodynamic size distributions of GO and purified GO:RGD in water.** Representative DLS size distributions of GO and GO:RGD are shown at 0, 4 and 24 h. Measurements were obtained from three independently prepared samples, each analysed in technical triplicate.

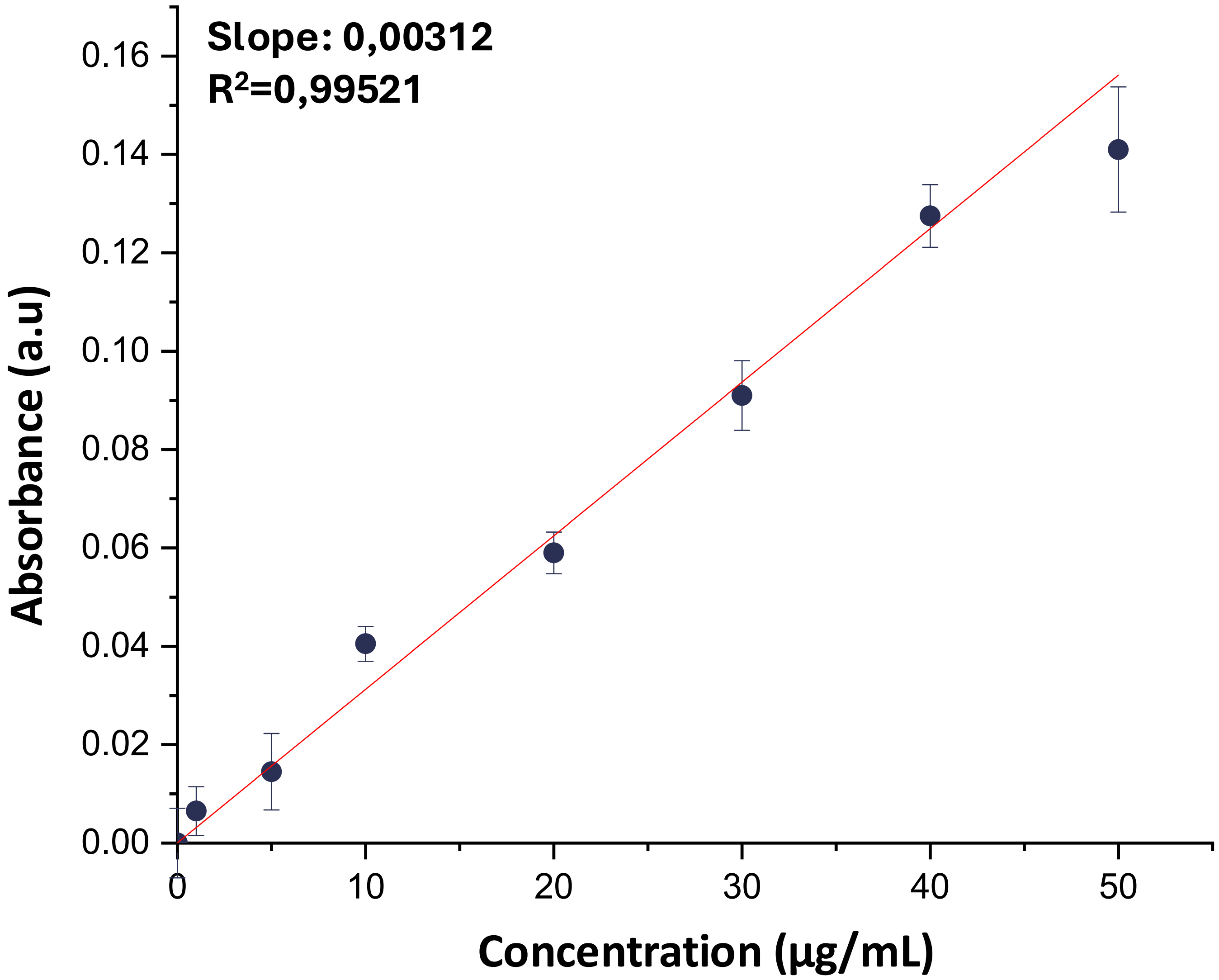


**Figure S2. RGD calibration curve for the TNBSA assay.** RGD standards ranging from 0.5 to 50 µg/mL were prepared in 0.1 M sodium bicarbonate solution and reacted with TNBSA, followed by absorbance measurement at 340 nm. Linear regression produced a slope of 0.00312 and $R^2$ = 0.99521. The calibration curve was used to calculate unbound RGD concentrations in the GO:RGD filtrates. Data are presented as mean ± SD of technical triplicate measurements.

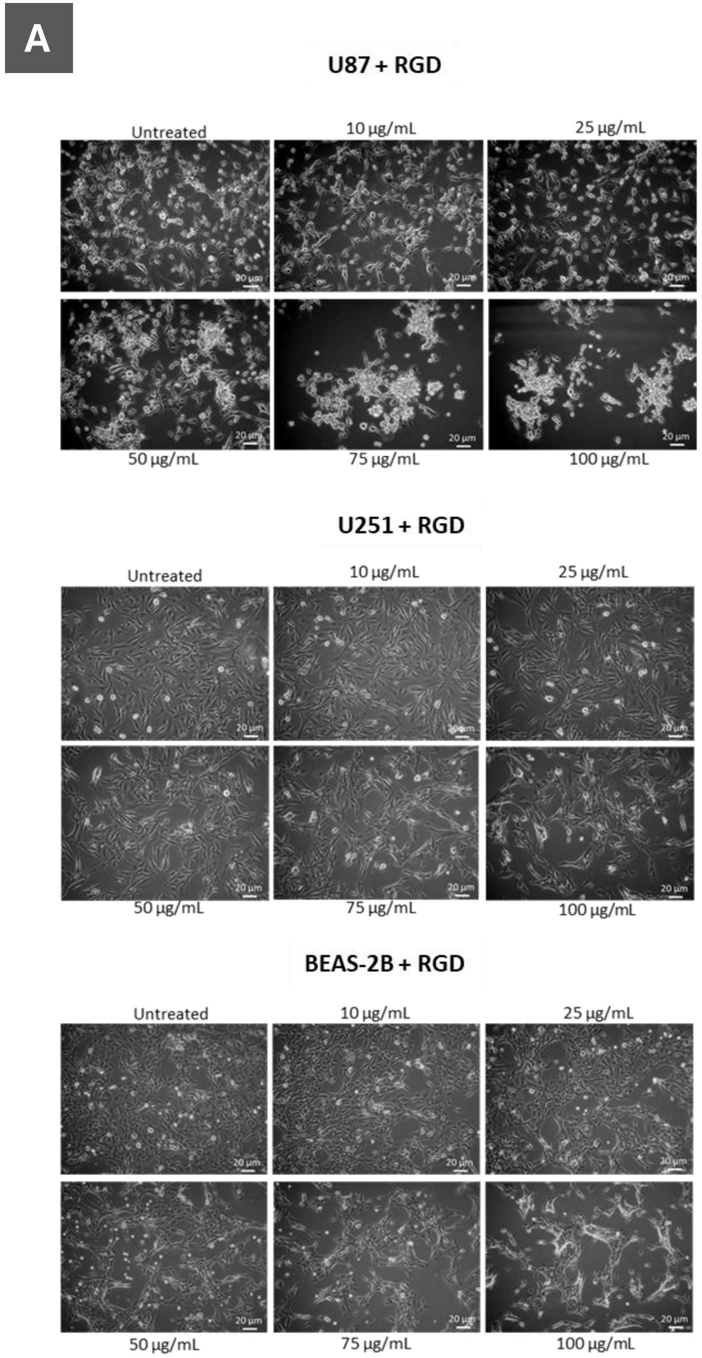


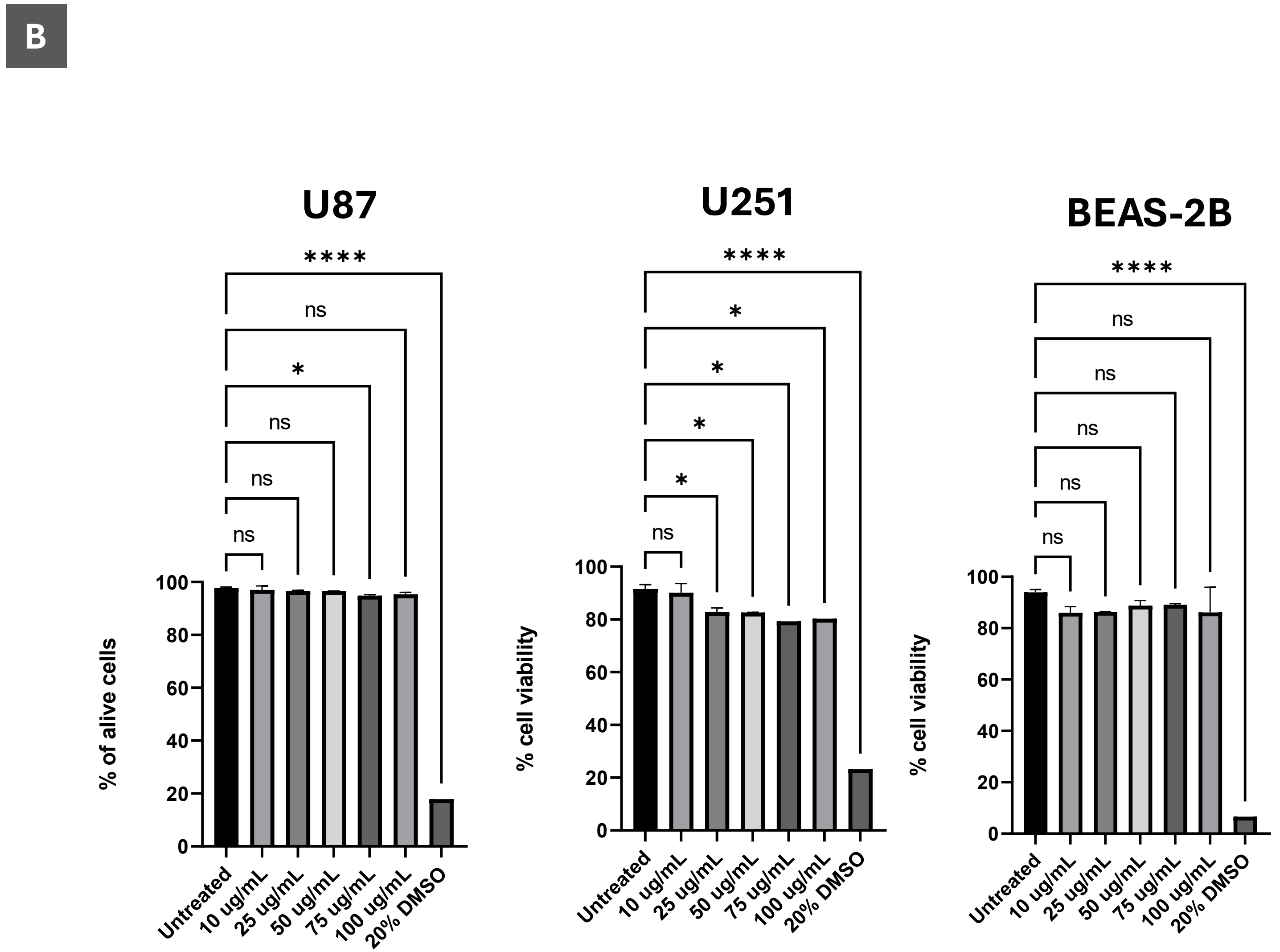


**Figure S3. Effect of free RGD on cell morphology and viability.** U87, U251 and BEAS-2B cells were exposed to RGD at 10, 25, 50, 75 or 100 µg/mL for 24 h. Untreated cells served as the negative control, and 20% DMSO served as the positive cytotoxicity control. (A) Representative bright-field micrographs showing cellular morphology following RGD exposure. Scale bars, 20 µm. (B) Cell viability determined by Annexin V/propidium iodide flow cytometry and expressed as the percentage of viable cells. Data are presented as mean ± SD from two independent experiments ($n = 2$), with duplicate measurements averaged within each experiment. Comparisons were performed separately for each cell line using one-way ANOVA followed by Dunnett's multiple-comparisons test against untreated cells. *$p < 0.05$, **$p < 0.01$, ***$p < 0.001$ and ****$p < 0.0001$; ns, not significant.

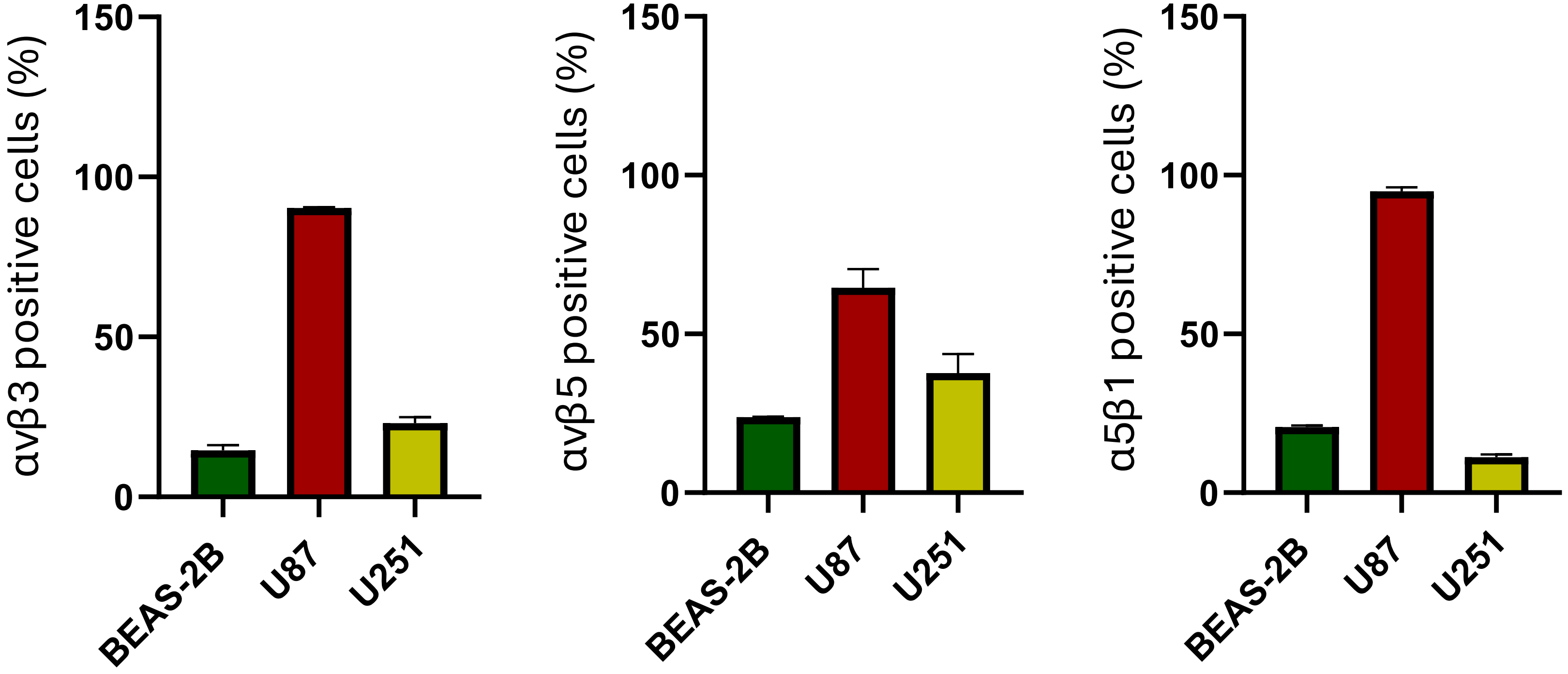


**Figure S4. Baseline proportions of αvβ3-, αvβ5- and α5β1-positive cells (%).** The percentages of U87, U251 and BEAS-2B cells positive for αvβ3, αvβ5 or α5β1 integrin were determined by flow cytometry using integrin-specific antibodies. Data are presented as mean ± SD from two independent experiments (n = 2), with technical triplicates averaged within each experiment.

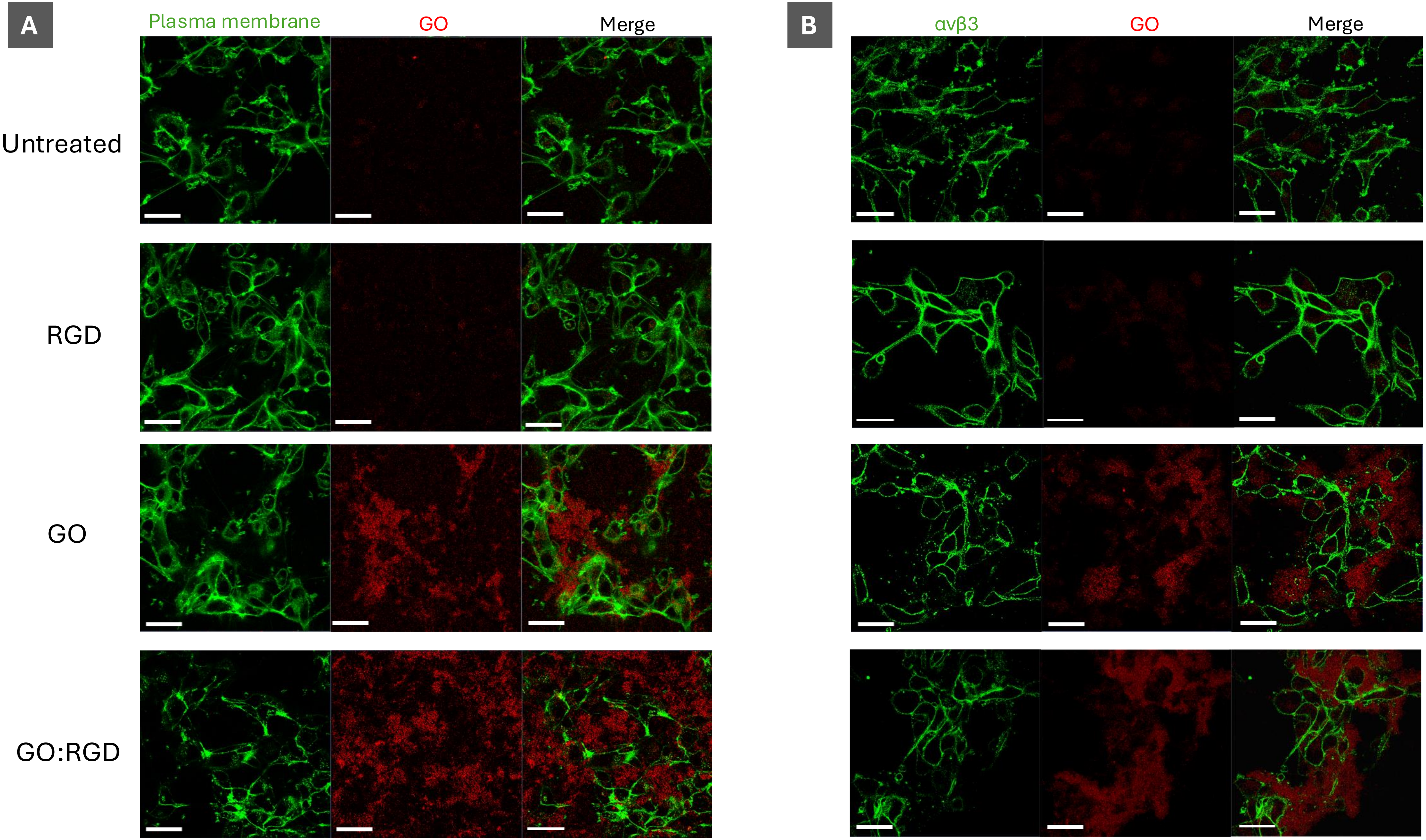


**Figure S5. Localisation of GO and GO:RGD relative to the plasma membrane and αvβ3-labelled regions in U87 cells.** Cells were untreated or exposed to RGD (5 μg/mL), GO (25 μg/mL) or GO:RGD (25:5 μg/mL) for 24 h. (A) The plasma membrane was labelled with CellMask Green. (B) αvβ3 was visualised by immunofluorescence staining in green. GO and GO:RGD were visualised through the intrinsic red fluorescence of GO. Representative individual channels and corresponding merged confocal images are shown. Scale bars, 40 μm.

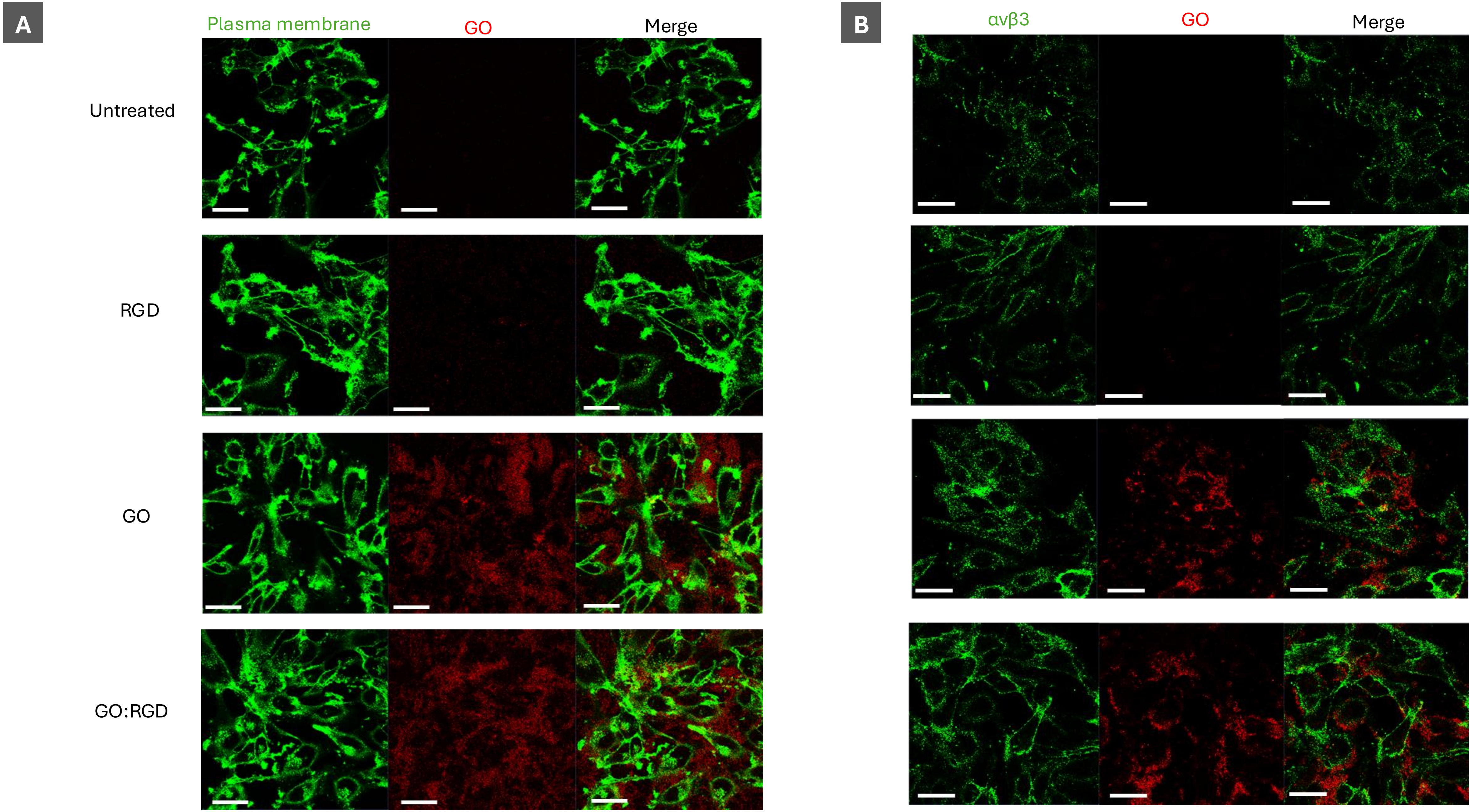


**Figure S6. Localisation of GO and GO:RGD relative to the plasma membrane and αvβ3-labelled regions in U251 cells.** Cells were untreated or exposed to RGD (5 μg/mL), GO (25 μg/mL) or GO:RGD (25:5 μg/mL) for 24 h. (A) The plasma membrane was labelled with CellMask Green. (B) αvβ3 was visualised by immunofluorescence staining in green. GO and GO:RGD were visualised through the intrinsic red fluorescence of GO. Representative individual channels and corresponding merged confocal images are shown. Scale bars, 40 μm.

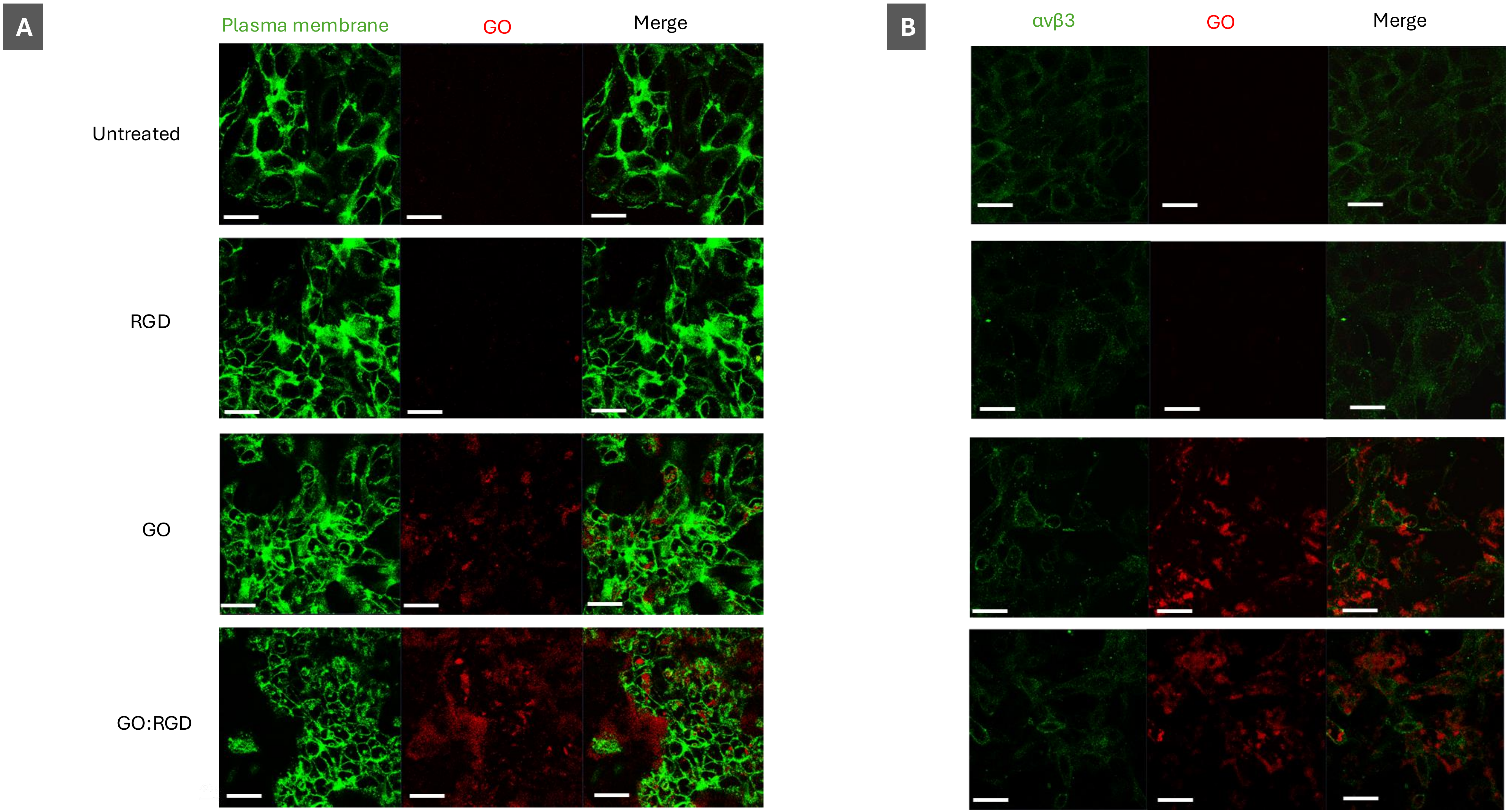


**Figure S7. Localisation of GO and GO:RGD relative to the plasma membrane and αvβ3-labelled regions in BEAS-2B cells.** Cells were untreated or exposed to RGD (5 μg/mL), GO (25 μg/mL) or GO:RGD (25:5 μg/mL) for 24 h. (A) The plasma membrane was labelled with CellMask Green. (B) αvβ3 was visualised by immunofluorescence staining in green. GO and GO:RGD were visualised through the intrinsic red fluorescence of GO. Representative individual channels and corresponding merged confocal images are shown. Scale bars, 40 μm.

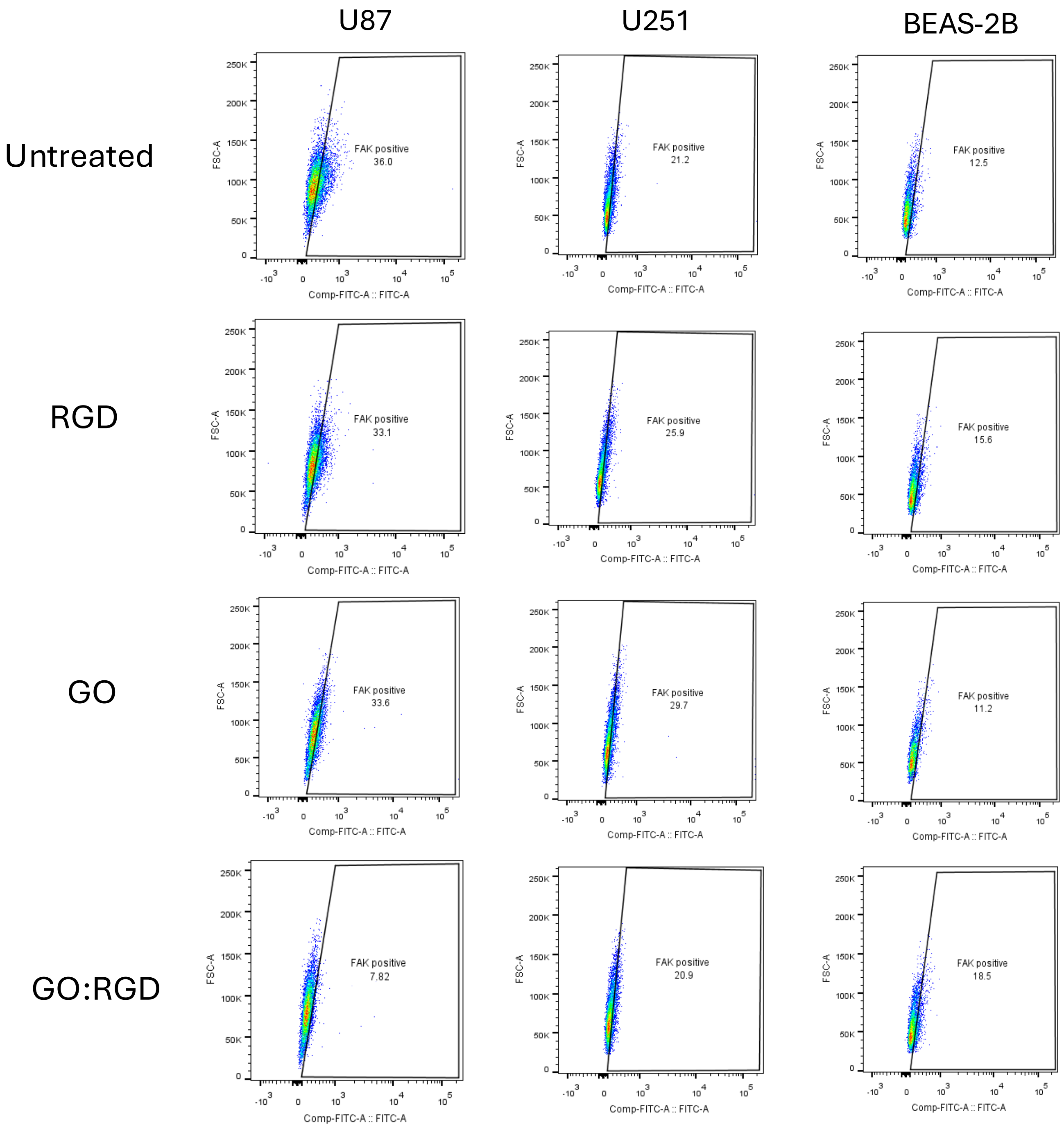


**Figure S8. Representative flow-cytometry plots showing the identification of pFAK Tyr397-positive cells.** U87, U251 and BEAS-2B cells were untreated or exposed to RGD (10 µg/mL), GO (50 µg/mL) or GO:RGD (50:10 µg/mL) for 24 h. Cells were fixed, permeabilised and stained for phosphorylated FAK at Tyr397. Rows correspond to the indicated treatments and columns to the three cell lines. The number shown beside each gate indicates the percentage of pFAK Tyr397-positive cells. Plots are representative of three independent experiments (n = 3), with duplicate measurements performed in each experiment.
